# Multidimensional Chromatin Regulation of Cell Lineage Differentiation in a Metazoan Embryo

**DOI:** 10.1101/2020.05.18.101584

**Authors:** Zhiguang Zhao, Rong Fan, Weina Xu, Yangyang Wang, Xuehua Ma, Zhuo Du

## Abstract

How chromatin dictates cell differentiation is an intriguing question in developmental biology. Here, a reporter gene integrated throughout the genome was used as a sensor to map the chromatin activity landscape in lineage-resolved cells during *C. elegans* embryogenesis. Single-cell analysis of chromatin dynamics across critical dimensions of cell differentiation was performed, including lineage, tissue, and symmetry. During lineage progression, chromatin gradually diversifies in general and exhibits switch-like changes following specific cell division, which is predictive of anterior-posterior fate asymmetry. Upon tissue differentiation, chromatin of cells from distinct lineages converge to tissue-specific states but retain “memory” of each cell’s lineage history, which contributes to intra-tissue heterogeneity. However, cells with a morphologically left-right symmetric organization utilize a predetermination chromatin strategy to program analogous regulatory states in early progenitor cells. Additionally, chromatin co-regulation drives the functional coordination of the genome. Collectively, this work reveals the role of multidimensional chromatin regulation in cell differentiation.

## INTRODUCTION

Chromatin state regulates diverse biological processes by controlling gene expression potential. During *in vivo* development, specific cells with identical genotypes differentiate into different cell types through the spatiotemporal expression of distinct sets of genes (Packer et al., 2019). Elucidating how dynamic and cell-specific chromatin states regulate *in vivo* cell differentiation processes has been a fundamental question in the fields of epigenetics and developmental biology (Yadav et al., 2018). Accurate assessments of chromatin state are essential to understanding how it functions in development. A combination of molecular approaches and high-throughput sequencing has been widely used to analyze the biochemical and biophysical properties of chromatin, including histone modifications, chromatin accessibility, and the spatial organization of chromatin (Goodwin et al., 2016). These epigenomic approaches have significantly enhanced the molecular profiling and mechanistic analysis of chromatin regulation during cell differentiation (Nicetto et al., 2019).

In addition to the epigenomic approaches described above, another powerful strategy for elucidating the functional state of chromatin is position-effect variegation (Elgin and Reuter, 2013; Schotta et al., 2003), which is a classic phenotype associated with *Drosophila* eye color. The white-eyed mutant phenotype is caused by the epigenetic silencing of normally active *white* genes, which is due to misplacement in heterochromatin regions (Timms et al., 2016). These findings established that chromatin environments exhibit strong positional effects on modulating the expression potential of nearby genes. The eye color phenotype provides a straightforward readout of the functional state of chromatin; thus, it has been widely used in genetic screens to identify potential regulators of chromatin activity. These efforts have identified many critical chromatin regulators that constitute much of our current knowledge of chromatin biology, including heterochromatin protein 1 (HP1) (James and Elgin, 1986), tri-methylation of histone H3 at lysine 9 (Rea et al., 2000), and histone deacetylase (Mottus et al., 2000). Furthermore, the positional effects on reporter gene expression have also been used to identify novel chromatin regulators in higher organisms, such as the human silencing hub complex HUSH (Ashe et al., 2008; Blewitt et al., 2005; Tchasovnikarova et al., 2015).

In addition to identifying chromatin regulators, positional effects on gene expression have also been exploited to study the different states of chromatin across the genome. In these studies, the expression status of the same reporter gene integrated into dozens to thousands of different genomic positions has been used as a chromatin activity sensor (Akhtar et al., 2013; Frokjaer-Jensen et al., 2016; Gierman et al., 2007). The analysis of positional effects across the genome has provided insight into several aspects of how the chromatin state is organized throughout the genome how it is regulated, including chromatin domain organization (Akhtar et al., 2013; Gierman et al., 2007), the structural properties of chromatin during the regulation of gene expression (Gierman et al., 2007), the repressive role of the lamina during gene expression (Akhtar et al., 2013) and germline gene silencing (Frokjaer-Jensen et al., 2016).

As compared to existing sequencing-based epigenomic approaches, the assessment of positional effects represents a unique approach that can be used to measure the functional state of chromatin as it relates to its role in regulating gene expression. However, applying this approach to developing single cells and using it to elucidate the regulatory roles played by chromatin during cell differentiation remains challenging. Previous large-scale analyses of positional effects primarily focused on dissecting the genomic properties and regulation of chromatin activity. Thus, these studies involved single-cell organisms, cell lines, and measuring reporter gene expression in multicellular organisms at the tissue/organism level. A systematic analysis of positional effects associated with single cells during *in vivo* differentiation has not been previously performed.

In this work, we exploited the positional effects in single cells to explore chromatin regulation of cell differentiation in *C. elegans*, a widely-used multicellular model organism in the field of single-cell developmental biology. Using a live-imaging approach, we quantified the expression levels of a reporter gene that was integrated into more than 100 positions throughout the genome and the resulting data were used to construct chromatin activity landscapes corresponding to lineage-resolved single cells during *C. elegans* embryogenesis. This analysis revealed the general dynamic patterns and chromatin regulatory strategies associated with critical processes of *in vivo* cell differentiation, including lineage commitment, anterior-posterior fate asymmetry, tissue differentiation, cell heterogeneity, and bilateral symmetry establishment. These findings will contribute to a systems-level mechanistic understanding of how chromatin regulates cell lineage differentiation in a metazoan embryo across cell lineages, tissue types, and symmetric morphological organization.

## RESULTS

### Quantification of the Positional Effects on Reporter Gene Expression in Lineage-Resolved Single Cells

In a previous study, a collection of *C. elegans* transgenic strains was generated, each containing the same GFP-expression cassette that had integrated into a distinct genomic position in single-copy (Figure 1A). The GFP is nucleus-localized and its expression is driven by the promoter of *eef-1A.1* (P*eef-1A.1*::GFP), a ubiquitously expressed translational elongation factor (Frokjaer-Jensen et al., 2014). Images of representative strains revealed strong position- and cell-dependent variation of GFP expression (Figure 1B), which suggests that P*eef-1A.1*::GFP is highly responsive to localized chromatin environments and that chromatin activity at many positions exhibits considerable cell specificity.

**Figure 1.**
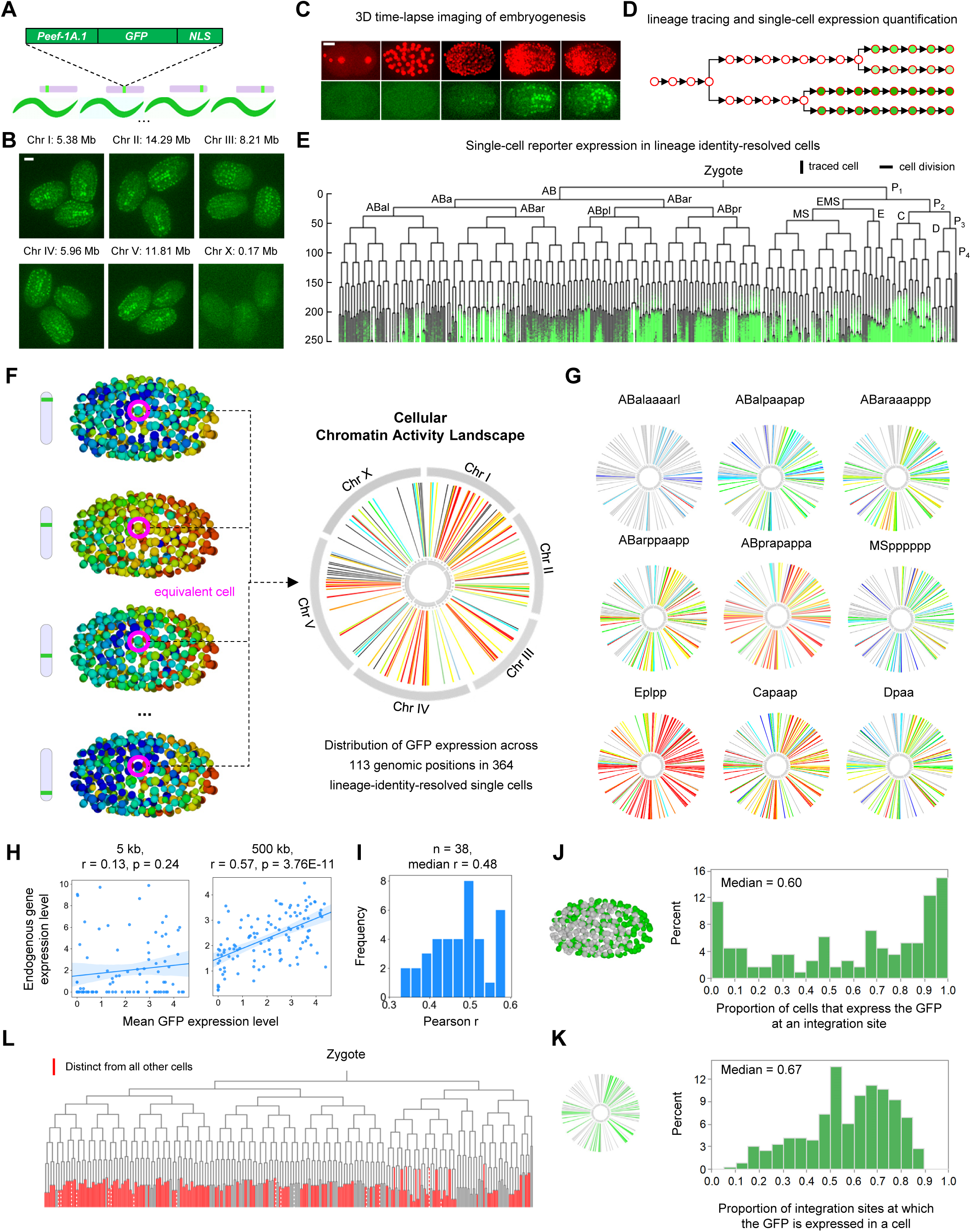
Quantification of Positional Effects on Reporter Expression in Lineage-Resolved Single Cells. (A) Transgenic strains carrying the same P*eef-1A.1*::GFP::NLS expression cassette integrated into different genomic positions. NLS denotes the nuclear localization signal. (B) Representative micrographs showing embryonic expression of GFP in representative strains. Scale bar = 10 µm. (C) 3D time-lapse imaging of embryogenesis. Scale bar = 5 µm. (D) Schematic of automated cell identification (circles), lineage tracing (arrows with bifurcations indicating cell divisions), and continuous quantification of GFP expression levels (green). (E) The color-coded tree shows single-cell expression of GFP (green) integrated into a position in each lineaged cell. Vertical lines indicate cells traced over time and horizontal lines indicate cell divisions. (F) Synthesis of cellular CAL. Left: the strategy used to synthesize CAL by integrating GFP expression across positions in equivalent cells (magenta). Right: The circos plot displays GFP expression levels at all genomic positions in a cell. (G) CAL of representative embryonic cells. (H) Correlation of expression levels between GFP and endogenous genes in 5-kb and 500-kb intervals centered on the integration sites. Statistics: Pearson correlation. (I) Distribution of Pearson correlation coefficient between single-cell expression levels of GFP and endogenous genes located in a 500-kb interval centered on the integration sites. Only the transcriptomes of cells that have been assigned with a unique identity were analyzed. Analysis of the transcriptomes of cells that have been assigned two possible identities yielded similar results. See also Table S2. (J) Distribution of the proportion of cells that express GFP at each integration site. (K) Distribution of the proportion of integration sites at which GFP is expressed in each cell. (L) Cells uniquely defined by CAL. Red indicates the cells in which the on/off status of GFP at five or more integration sites was distinct from other cells. See also Figures S1, S2, and Table S2.

To facilitate single-cell quantification of GFP expression, we generated a collection of dual-fluorescent strains by introducing another nucleus-localized, ubiquitously expressed mCherry transgene into the above strains for nuclei identification and tracing (Table S1). We used 3D time-lapse imaging to record embryogenesis at high spatiotemporal resolution and images were analyzed to determine GFP expression in individual cells (Du et al., 2014; Murray et al., 2008) (Figures 1C). Cell lineages were reconstructed based on the automatic identification and tracing of all cells via the mCherry signal, followed by multiple rounds of manual curations (Figure 1D). Simultaneously, the intensity of GFP in each traced nucleus was measured to indicate chromatin activity at a certain genomic position (Figure 1E). In total, positional effects on GFP expression was quantified at 113 genomic positions (0.88 Mb genomic resolution) in 268 embryos (Table S2). Each of the processed embryos yielded quantitative GFP expression in 727 lineaged cells covering all cells until the 350-cell stage, when the embryo has completed tissue fate determination (Sulston et al., 1983).

We validated the accuracy of the annotation of GFP integration sites that had been done previously and the reliability of the lineage tracing results (STAR Methods; Figure S1A). Additionally, we optimized the expression quantification method to compensate for the attenuation of fluorescence intensity with sample depth and to precisely measure the instantaneous GFP expression in a single cell (Figure S1B-E). These improvements allowed for a more reliable comparison of inter-cell gene expression. Cellular GFP expression was highly reproducible, with a median correlation coefficient of 0.83 and a median consistency of binarized expression of 0.90 between experimental replicates (Figure S1F). Aggregating GFP expression across all positions showed that P*eef-1A.1*::GFP was predominantly expressed in the majority of cells from the 350-cell stage (with 364 cells) and onward (Figure S1G), making the analysis of chromatin activity in cells from this stage the most informative. Unless otherwise specified, all analyses focused on the 364 traced terminal cells.

### Cellular Chromatin Activity Landscape is Highly Dynamic and Informative for Defining Cellular States

*C. elegans* embryogenesis follows a fixed cell lineage to generate the same set of terminally differentiated cells that are identical and comparable between embryos at single-cell resolution (Sulston et al., 1983). Although cellular GFP expression at each integration site was assayed in individual embryos, the invariant nature of the cell lineage allowed precise integration of expression levels at different positions of the genome in lineage-equivalent cells (Figure S2A and Figure 1F). Using GFP expression as an indicator of chromatin activity, we constructed the distribution of chromatin activity across 113 genomic positions (termed chromatin activity landscape, CAL) for all 364 somatic cells at the 350-cell stage (Figure 1G and Table S2).

Both bulk and single-cell analyses confirmed that the measured positional effects on P*eef-1A.1*::GFP reliably indicated chromatin activity. Based on the bulk analysis, GFP expression levels were highly consistent with various properties of chromatin closely related to its activity, including genome distribution, histone modification pattern, chromatin accessibility, and the 3D organization of chromatin (Figure S2B-F and Table S2; STAR Methods). Conversely, GFP expression was not significantly associated with local genetic environments. Integration of the transgene into intragenic and intergenic regions resulted in comparable expression levels (Figure S2G). Furthermore, no correlation between the expression level of GFP and endogenous genes located in a 5-kb interval centered on the integration sites was observed (Figure 1H, left, r = 0.13, p = 0.24), indicating that local genetic environments did not significantly affect GFP expression. However, a strong correlation was observed when the size of the interval was extended to over 100 kb, with the 500-kb interval yielding the strongest correlation (Figures 1H, right, r = 0.57, p = 3.76E-11). Since gene expression over a large chromosome domain is more likely to be governed by chromatin activity, this result again suggests that the positional effects on GFP expression are due to chromatin activity. Single-cell analysis also supported the interpretation that the positional effects on GFP expression are indicative of chromatin activity. Recently generated lineage-resolved single-cell RNA-sequencing (scRNA-seq) data of *C. elegans* embryogenesis (Packer et al., 2019) confirmed the correlation between the expression levels of GFP and an endogenous gene over a large chromatin domain in single cells (r = 0.48, Figure 1I). However, due to the scarcity of single-cell chromatin data, we were unable to assess the relevance of cellular chromatin activity in the context of other chromatin properties.

Intuitively, the CAL was highly dynamic across integration sites and between cells (Figure 1G). Qualitatively, at each integration site, GFP was expressed in a proportion of the cells (median = 60%) (Figure 1J), which suggests that chromatin activity at most genomic positions was cell-specific. At only seven positions, GFP was constitutively active (n = 1) or silenced (n = 6). In each cell, GFP was expressed only when it had been integrated into a fraction (median = 67%) of genomic positions, suggesting that chromatin activity in a cell is positionally specific (Figure 1K). In no cells did we observe constitutive GFP expression (>90%) at all integration sites, suggesting that *eef-1A.1* promoter activity is not cell- or tissue-specific (if it were, then a considerable number of cells would constitutively express GFP).

To address whether the specific CAL can distinguish individual cells, we calculated the proportion of cells that can be uniquely defined by CAL, which showed that, intriguingly, the CAL uniquely defined as many as 67% of the cells. In these cells, the on/off state of chromatin activity at five or more genomic positions was distinct from that of all other cells (Figure 1L). Such a fraction is substantial because many embryonic cells share a similar developmental fate and differentiate into the same tissue type. Thus, the CAL provides rich information that can be used to distinguish cells at a sub-tissue level.

In summary, the data presented above systematically confirmed that cellular CAL is highly dynamic, informative, and biologically relevant. With lineage-resolved CAL and extensive prior knowledge of the developmental function of each *C. elegans* cell, we next systematically investigated chromatin regulation of cell differentiation, as described in the following sections. Specifically, the cellular dynamics of CAL were used to elucidate the developmental role of chromatin in critical regulatory processes of cell differentiation, including lineage specification, anterior-posterior fate asymmetry, tissue differentiation, cell heterogeneity, and bilateral symmetry establishment.

### Gradual Diversification of CAL Recapitulates Global Kinetics of Lineage-Coupled Cell Differentiation

Cell differentiation accompanies lineage progression. The lineage-based mechanism plays a crucial role in initiating cell differentiation in the early embryo by assigning developmental fates to progenitor cells in a lineage-identity-dependent manner (Labouesse and Mango, 1999). Changes in CAL across cell lineages and whether it can predict the kinetics of lineage-coupled cell differentiation were evaluated.

We first quantified CAL divergence as a function of the lineage relationship between 364 traced terminal cells. The lineage relationship was quantified as cell lineage distance, which was calculated as the total number of cell divisions separating the cells from their lowest common ancestor (Figure S3A). In the majority of cases, a higher CAL divergence was observed between cells with a large lineage distance and, globally, CAL divergences increased progressively with cell lineage distance (Figures 2A, 2B, and Table S3). Thus, in general, CAL diversifies gradually across cells during lineage progression.

**Figure 2.**
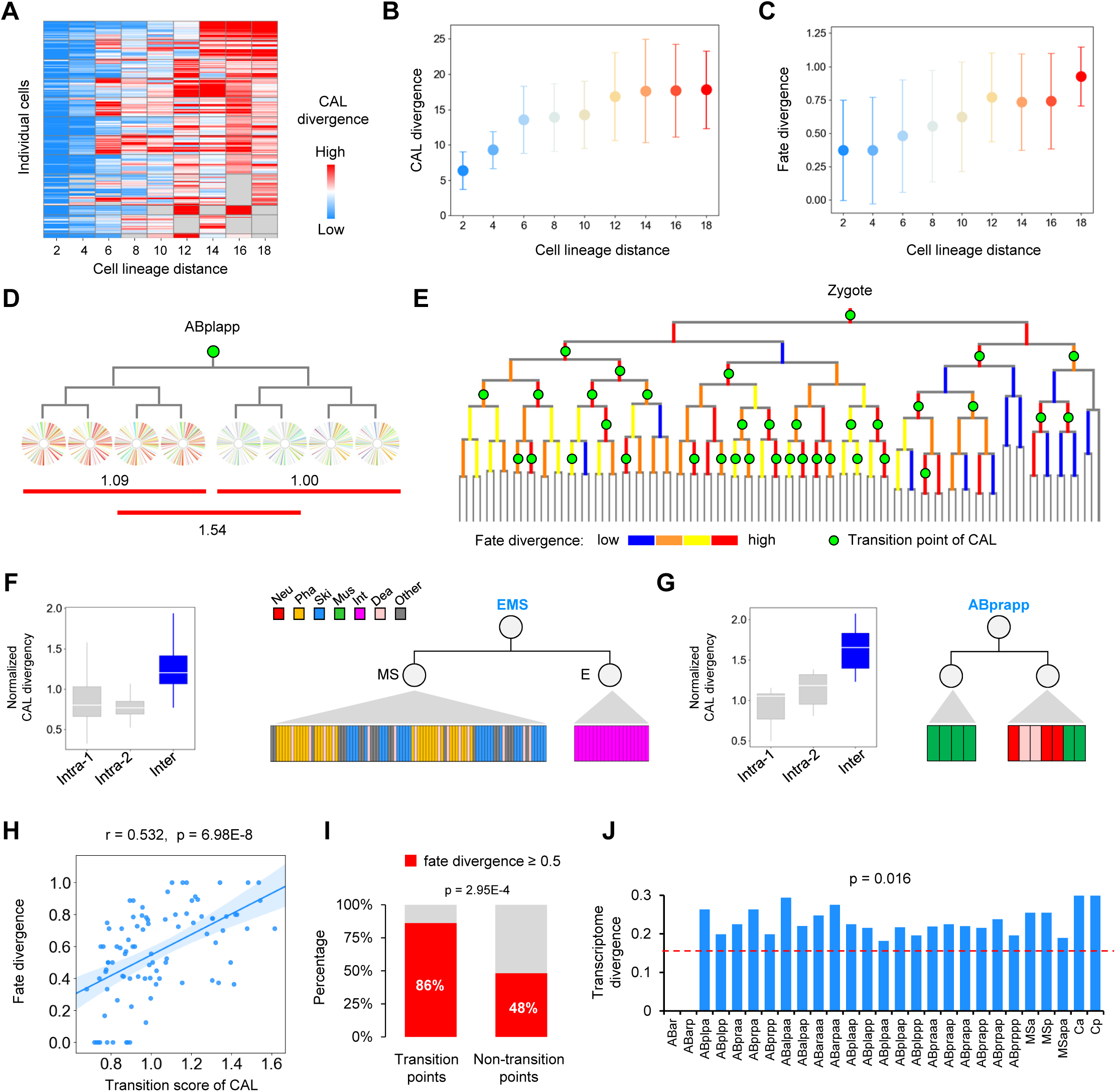
CAL Dynamics During Lineage Progression Informs Lineage-Coupled Cell Differentiation. (A) Heatmap showing the mean CAL divergence between each of the 364 traced terminal cells to all other cells at different cell lineage distances. (B) Changes in CAL divergence following an increase in cell lineage distance. Because an odd-numbered lineage distance involves cells at different generations, only those with an even number (80% of the cases) were used to analyze the relationship between cell lineage distance and other cellular attributes. Data are presented as mean ± SD. (C) Changes in fate divergence following an increase in cell lineage distance at characteristic developmental stages. Data are presented as mean ± SD. (D) Representative CAL transition point. (E) Tree visualization of the inferred CAL transition points (green circles) and associated fate divergence between daughter cells of progenitor cells (color-coded vertical lines). (F-G) Two examples showing the association between CAL transition and anterior-posterior fate asymmetry following EMS (F) and ABprapp (G) cell division. Left: the intra- and inter-daughter-lineage CAL divergences, Right: the fates of two daughter cells following the division. (H) Correlation between CAL transition and fate divergence in early progenitor cells (n = 90). (I) Comparison of the percentage of high fate divergence (≥0.5) between CAL transition and non-transition points. Statistics: Fisher’s exact test. (J) Transcriptome divergence between daughter cells following a cell division that exhibits CAL transition. The dashed line indicates the mean transcriptome divergence between daughter cells following a non-CAL-transition cell division. Statistics: Mann-Whitney U test. See also Figure S3 and Table S3.

We next analyzed the lineage-coupled kinetics of cell differentiation by measuring how the cell fates change as a function of the lineage distance between cells. Based on the lineage tree structure and tissue types of all terminally differentiated cells, we retrospectively defined the fate of each cell as the combinatorial pattern of tissue types produced by each cell and quantified the fate difference between cells (STAR Methods and Figure S3B). This analysis showed that as the cell lineage unfolds, cells differentiate progressively, in general. The fate divergence between cells was proportional to their lineage distances at different developmental stages, recapitulated that of CAL dynamics (Figure 2C and Figure S3C). To further demonstrate that CAL dynamics were associated with fate changes, the relationship between the two was directly analyzed between cells. The result showed that a higher CAL divergence was generally associated with a higher fate divergence, especially between cells at a modest lineage distance (from 6 to 14) (Figure S3D). Thus, CAL dynamics during lineage progression recapitulate the kinetics of cell differentiation.

### CAL Transitions Predict Anterior-Posterior Fate Asymmetry

An important paradigm of the lineage-based mechanism for regulating cell differentiation is asymmetric cell division, in which a mother cell generates two daughter cells with distinct developmental fates (Mizumoto and Sawa, 2007). Since the majority of *C. elegans* cell divisions occur along the anterior-posterior body axis, this asymmetry is also termed anterior-posterior fate asymmetry. For example, the progenitor cell named EMS produces two daughter cells with distinct developmental fates: the anterior daughter MS differentiates into mesodermal tissues and the posterior daughter E differentiates into endodermal tissue (Thorpe et al., 1997).

We determined whether CAL changes in a switch-like fashion following specific cell divisions and whether this is indicative of anterior-posterior fate asymmetry. For this, we used a retrospective strategy to infer the CAL of a progenitor cell by the CAL of all terminal cells the progenitor cell generates during development. This assumption is based on the parsimony principle, which has been widely used in biology in which a minimal set of regulatory events are used to explain the observed outcomes. A cell division is defined as a transition point of CAL if CAL divergences between terminal cells generated by different daughter cells (inter-divergence) are significantly higher than those between terminal cells generated by the same daughter cell (intra-divergence) (STAR Methods, Figure S3E and Table S3). As exemplified by Figure 2D, division of the ABplapp cell is identified as a CAL transition point, since cells generated by its two daughter cells exhibit distinct CAL. Of 90 early cell divisions, 36 (40%) of them showed significant CAL transitions (Figure 2E). Within these transition points, we correctly captured the division of the EMS cell as being a CAL transition point, which is consistent with the knowledge that Wnt signaling induces an anterior-posterior fate asymmetry (Figure 2F) (Thorpe et al., 1997). We also identified CAL transition points in many cases in which the cell division produces two daughter cells with distinct developmental fates. For example, a CAL transition point was identified during asymmetric cell division of the ABalpap cell that produces an anterior daughter that exclusively differentiated into skin cells and a posterior daughter that differentiated into skin and neuronal cells and cells that undergo programmed cell death (Figure 2G). Globally, the CAL transition score correlated significantly with the fate divergence between two daughter cells (Figure 2H, r = 0.595, p = 6.25E-10). Of the identified transition points of CAL, 86% of them had concomitant high fate divergence (≥ 0.5), which was significantly higher than what was observed in non-transition points (Figure 2I). Thus, CAL transitions that followed specific cell divisions are predictive of anterior-posterior fate asymmetry.

Since the above analyses were based on a retrospective approach, it is unclear whether the inferred transition of CAL occurred in the daughter cells or later during development. Using the scRNA-seq dataset (Packer et al., 2019), we directly compared transcriptome divergence between two daughter cells and used it as a proxy for cell fate/state. Consistent with our results, significantly larger transcriptome divergences between daughter cells were observed following the cell division that exhibited a CAL transition as compared to those following other cell divisions (Figure 2J). This result confirmed that CAL transitions are predictive of anterior-posterior fate asymmetry.

### Characteristic CAL is Established in Specific Cell Lineages

The direct outcome of lineage-based regulation of cell differentiation is the assignment of specific developmental fates to specific progenitor cells according to their lineage identity. For example, the ABalp progenitor cell is always assigned the developmental fate of generating pharyngeal and neuronal cells, whereas the E progenitor cell invariantly is assigned the fate of differentiating into intestinal cells.

First, we determined to what extent characteristic CALs are established in cells from different lineages. We divided the entire cell lineage into smaller lineage groups and compared chromatin activity across all genomic positions between cells from different lineage groups. We found CAL differed in cells from different lineage groups (Figure 3A), based on quantification of the fraction of genomic positions at which the on/off state of chromatin activity was different between two cells. For example, after dividing the cell lineage into 12 groups, chromatin activity at 20% of all positions (ranging between 6% to 42%) of cells from ABala lineages was different, on average, as compared to all other lineages. Thus, characteristic CAL was established in different lineages, which accompanied the process of lineage fate specification.

**Figure 3.**
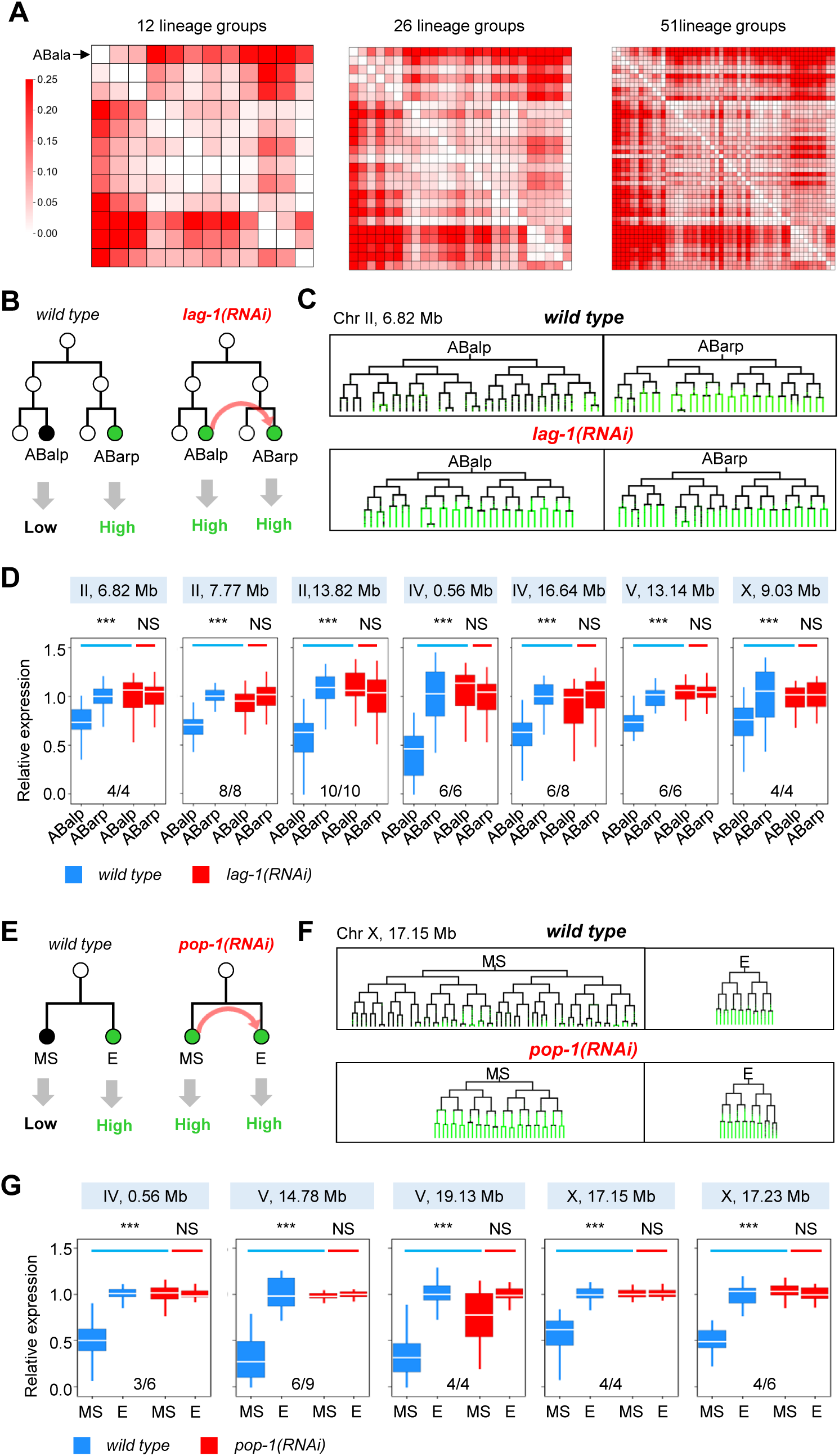
Characteristic CAL is Established in Specific Cell Lineages According to Developmental Fates. (A) Heatmap showing the average fraction of genomic positions exhibiting distinct on/off states of chromatin activity between cells in all pair-wise lineage comparisons. The value from intra-lineage comparison was subtracted from the value of the corresponding inter-lineage comparison. (B) Top: Schematic of ABalp-to-ABarp lineage fate transformation (red arrow) induced by *lag-1(RNAi).* Bottom: predicted changes in chromatin activity in cells from corresponding lineages. (C) Example of GFP expression in cells from the ABalp and ABarp lineages before and after lineage fate transformation. (D) Each panel shows the relative expression levels of GFP at an integration site in cells from the ABalp and ABarp lineages in wild-type and *lag-1(RNAi)* embryos. GFP expression levels of all cells were normalized to the mean expression levels of cells from the ABarp lineage. Multiple embryos were analyzed for each genotype and the ratio indicates how many times (wild type-RNAi comparisons) the changes in GFP expression are consistent with the expectation. Statistics: Mann-Whitney U test (***, p < 0.001, NS, p > 0.05). (E) Top: Schematic of MS-to-E lineage fate transformation induced by *pop-1(RNAi).* Bottom: the predicted changes in chromatin activity in cells from corresponding lineages. (F) An example of GFP expression in cells from the MS and E lineages before and after lineage fate transformation. (G) Changes of GFP expression upon MS-to-E lineage fate transformation induced by *pop-1(RNAi)*. The organization of the figure is the same as in (C). GFP expression levels of all cells were normalized to the mean expression levels of cells from the E lineage. Statistics: Mann-Whitney U test (***, p < 0.001, NS, p > 0.05). See also Figure S4.

Next, we performed lineage fate perturbation experiments (STAR Methods) to test whether cellular CAL changes accordingly once lineage fates are switched. We first used the ABalp and ABarp lineages to address this question because cells derived from the two lineages exhibited significantly different CAL. RNA interference (RNAi) was used to knock down the function of the Notch effector *lag-1/CLS* (Moskowitz and Rothman, 1996), which induced an ABalp-to-ABarp lineage fate transformation (Figure 3B). In *lag-1(RNAi)* embryos, the tree topology and characteristic programmed cell death of the ABalp lineage were distinct from normal ABalp but resembled those of the ABarp lineage (Figures S4A and S4B), which confirmed the induction of lineage fate transformation. Concomitantly, in all seven integration strains that were examined, GFP, which is usually silenced or weakly expressed in ABalp lineage-derived cells, was highly expressed, similar to the normal ABarp lineage (Figures 3C and 3D and Figure S4C). These results suggest that CAL and cell lineage fate are coupled.

This coupling was confirmed in another developmental context. The fate of the MS lineage was switched to that of E by targeting RNAi against the *pop-1* gene (Lin et al., 1995), a Wnt signaling component (Figures 3E and S4D). Concomitantly, in all five tested integration strains, GFP was up-regulated in MS lineage-derived cells, resembling that of typical E lineage-derived cells (Figures 3F, 3G, and S4E).

Collectively, these lineage-centric analyses revealed the existence of a lineage-based chromatin regulation of cell differentiation. Globally, CAL diversifies gradually following lineage progression, which recapitulates the kinetics of cell differentiation. In certain contexts, cellular CAL changes in a switch-like fashion between daughter cells that is predictive of anterior-posterior fate asymmetry. Finally, characteristic CALs are established in specific lineages, which are tightly coupled to their development fates.

### A Convergence Chromatin Strategy to Drive Tissue Differentiation in Cells From Multiple Lineages

The lineage-based mechanism archives a global patterning of cell fates in groups of lineage-related cells. Since most cell types are not monoclonal, this mechanism alone is not sufficient to generate the ultimate cell types required by an organism. Complementarily, there is a tissue-based mechanism that refines cell differentiation, whereby the same tissue fate is co-specified in multiple tissue precursors (Labouesse and Mango, 1999). This tissue-based mechanism is involved throughout embryogenesis because most somatic tissues, except the intestine, are derived from multiple cell lineages (Figure 4A).

**Figure 4.**
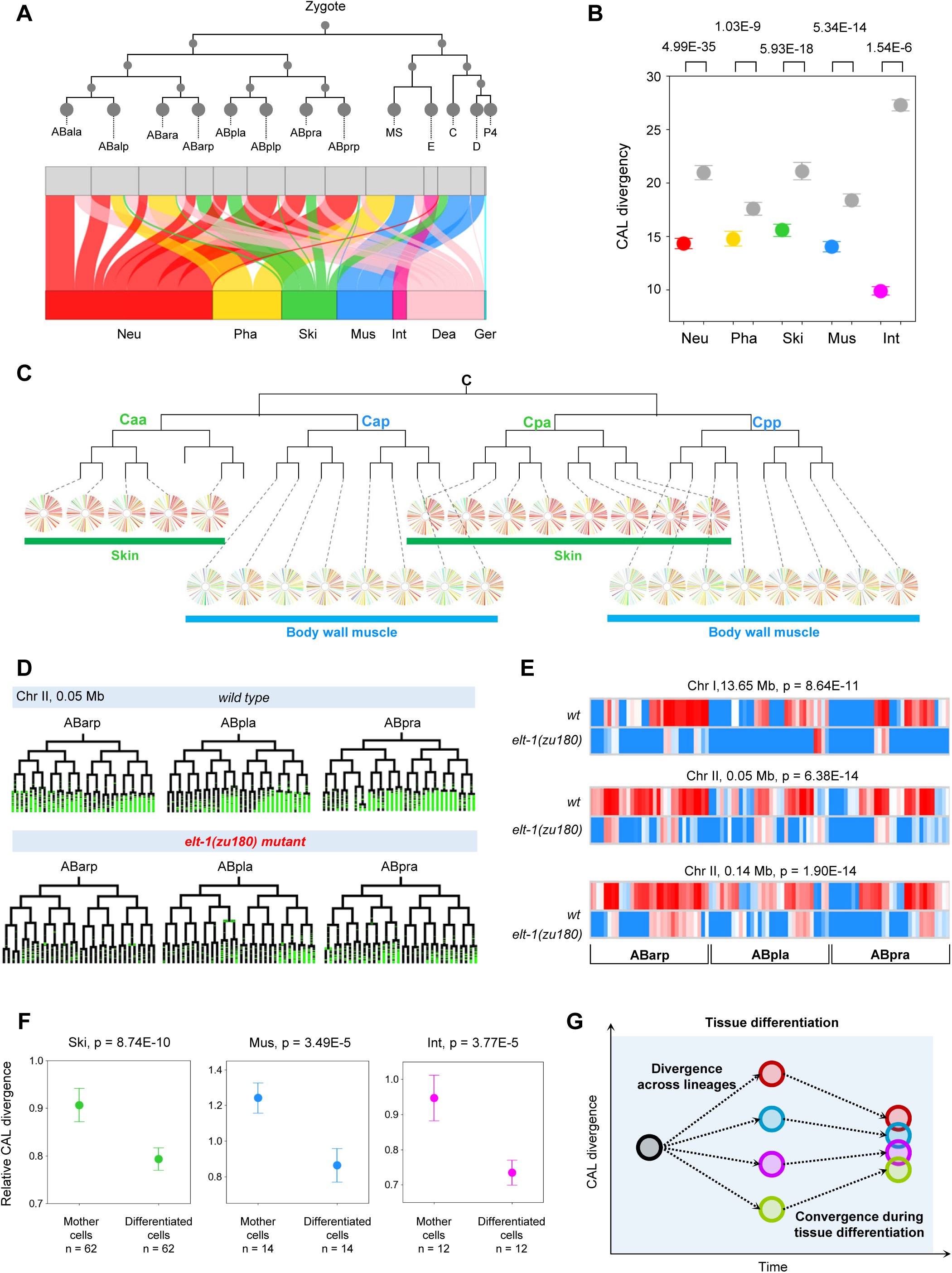
Convergence of CAL Accompanies Tissue Fate Differentiation. (A) The relationship between cell lineage identities (12 somatic founder cells) and tissue fate. Neu, neuronal system; Pha, pharynx; Ski, skin; Mus, body wall muscle; Int, intestine; Dea, cell death; Ger, germ cell. (B) CAL divergence between intra- and inter-tissue cells for each tissue. Data are presented as mean ± 95% CI. Statistics: Mann-Whitney U test (C) An example showing the convergence of CAL in skin and body wall muscle cells from the C lineage. (D) An example showing changes in GFP expression in cells from the ABarp, ABpla, and ABpra lineages before and after skin fate perturbation. (E) Expression levels of GFP at three integration sites in individual cells from three major skin cell lineages in wild-type and *elt-1(zu180)* embryos. Statistics: Wilcoxon signed-rank test. See Figure S5 for additional embryos. (F) CAL divergence between differentiated intra-tissue cells and between their mother cells for each tissue. The neuronal system and pharynx are not included because of a small number of differentiated cells at the 350-cell stage. Data are presented as mean ± 95% CI. Statistics: Mann-Whitney U test. (G) The convergence chromatin strategy to regulate tissue fate differentiation. See the main text. See also Figure S5 and Table S4.

As shown above, CAL is generally diversified across cells from different lineages, which raises the question as to whether the CAL of cells from distinct lineages converges according to tissue types. If it does, then CAL divergence would be lower between cells of the same tissue (intra-tissue) as compared to between different tissues (inter-tissue). We classified all cells into five major tissue/organ types and found that, in all tissue types, intra-tissue CAL divergence was significantly lower as compared to inter-tissue (Figure 4B and Table S4). For example, although with distinct lineage origins, CAL of skin cells from the Caa and Cpa lineages was highly similar, so was CAL of body muscle cells derived from the Cap and Cpp lineages (Figure 4C).

Tissue fate perturbation experiments were performed to determine whether CAL is coupled to tissue fate. Specifically, a mutant of the *elt-1*/GATA1 gene, which is a specifier of skin fate (Page et al., 1997), was used to abolish skin differentiation. Three integration strains were used to profile chromatin activity. In *elt-1* mutant embryos, GFP expression levels in cells from the three major cell lineages (ABarp, ABpla, and ABpra) that normally differentiate into skin were significantly different as compared to wild-type embryos (Figures 4D, 4E, and S5A). These results confirm that characteristic CAL is established in specific tissues and is coupled to tissue fate.

The fact that CAL diversifies across cells in different lineages and that the CAL of cells belonging to the same tissue originated from different lineages are similar suggests a convergence of CAL during tissue differentiation. We found that CAL divergence between differentiated intra-tissue cells was significantly lower than the CAL divergence between their mother cells (Figure 4F), which suggests that convergence is progressive toward terminal tissue differentiation. Furthermore, progressive convergence was also supported by an analysis of cellular transcriptomes. Using the scRNA-seq data, we quantified transcriptome divergence between precursors of intra-tissue cells at different developmental stages. The results showed that divergence generally decreased from the undifferentiated stages to the terminally differentiated stages along the developmental trajectories of all tissue types (Figure S5B).

In summary, these results revealed a convergence chromatin strategy that specifies similar and characteristic regulatory states in cells from multiple lineages in a tissue-dependent manner.

### Chromatin Exhibits “Memory” of Lineage Origins that Contributes to Cell Heterogeneity

While the convergence of CAL generally specifies similar CAL in cells of the same tissue type, the extent of convergence was variable across tissues (Figure 5A). For example, the heterogeneity of CAL between neuronal cells was significantly higher than it was between intestinal cells. Furthermore, CAL heterogeneity between some cells was notably higher than that of other cells within the same tissue (Figure 5A). Given that CAL divergence increased with cell lineage distance (Figure 2B) and that cells of different tissues have significantly different lineage compositions (Figure 4A), we tested whether lineage origin of cells accounts for this heterogeneity. First, we found the CAL heterogeneity in each tissue was highly predictive by the lineage composition of cells. Tissue cells from very distant lineages tend to exhibit higher CAL heterogeneity; the correlation coefficient between the average cell lineage distance and intra-tissue CAL divergence was 0.98 (Figure 5B). Second, to further confirm these lineage effects, CAL divergence between intra-tissue cells was quantified as a function of cell lineage distance in each tissue, which showed that a larger lineage distance was generally associated with higher CAL divergence (Figure 5C). As exemplified by Figure 5D, CALs of skin cells that are more lineage related are more similar as compared to those with a more distant lineage relationship. These results demonstrate that the lineage origin of cells contributes to intra-tissue CAL heterogeneity.

**Figure 5.**
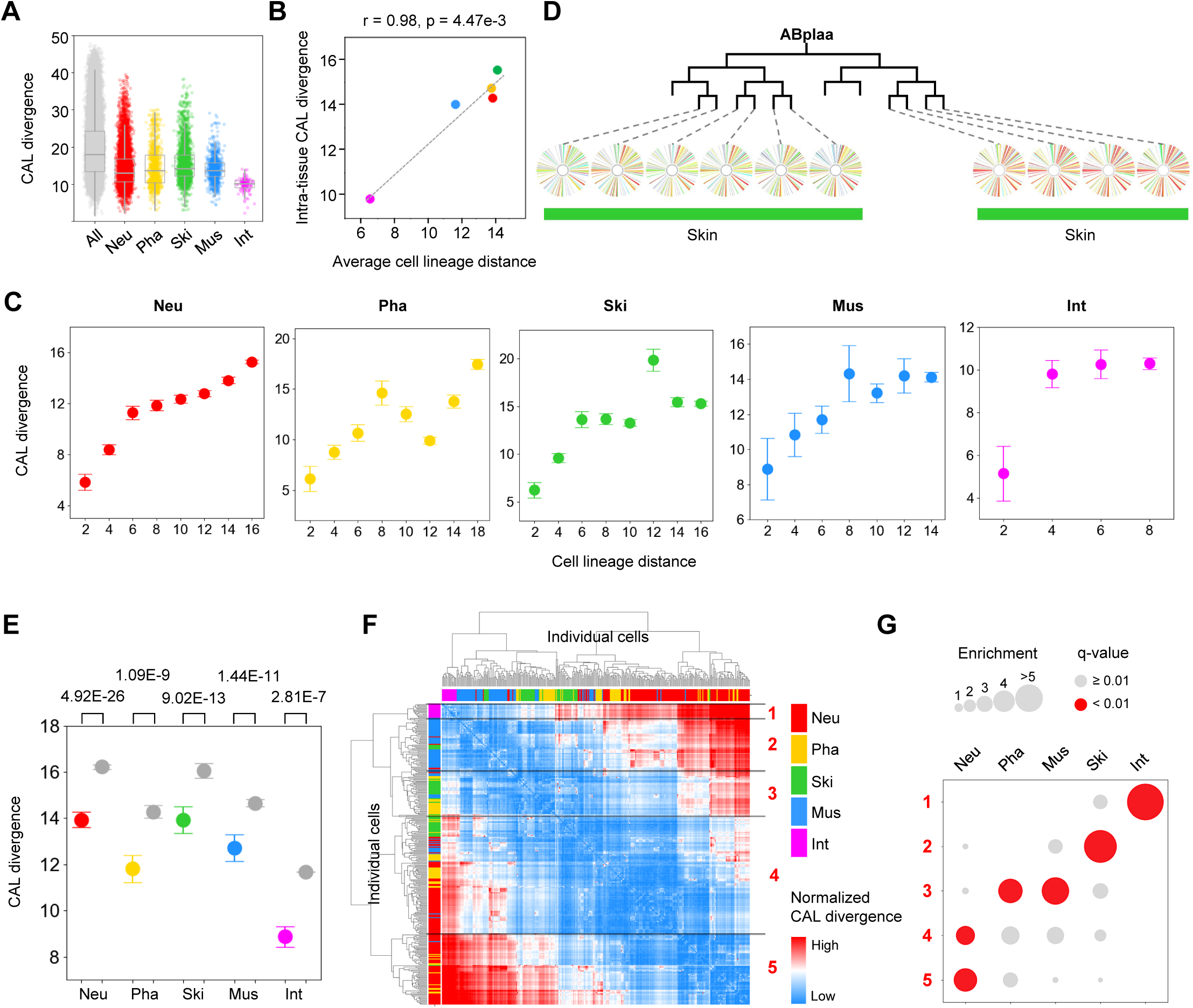
Lineage Origin Contribute to Intra-Tissue Heterogeneity. (A) Pair-wise comparison of CAL divergence between all cells and intra-tissue cells. (B) Correlation between average lineage distance and CAL divergence between intra-tissue cells for all tissues. (C) Changes in CAL divergence following the increase of cell lineage distance between intra-tissue cells. Data are presented as mean ± 95% CI. (D) An example showing that CAL of skin cells originated from different cell lineages. (E) CAL divergences of intra-tissue cells and inter-tissue cells (gray) after controlling for the influence of lineage distance between cells. For each pair of intra-tissue cells, the mean CAL divergence of all inter-tissue cells with identical cell lineage distance was used as the control. Data are presented as mean ± 95% CI. Statistics: Mann-Whitney U test. (F) Hierarchical clustering of cells using lineage distance normalized CAL divergences. (G) Enrichment of tissue types in cells in each cluster. Statistics: Fisher’s exact test, Benjamini-Hochberg adjusted P-value. See also Figure S6 and Table S5.

Not all of the 364 traced terminal cells had completed cell differentiation, which makes it unclear whether such effects were stably maintained in the CAL of terminally differentiated cells. We repeated the above analysis focusing on only post-mitotic cells within the traced cells, which are likely to have completed terminal differentiation (Table S4). A similar result was obtained (Figure S6A), showing that the lineage effects on intra-tissue chromatin heterogeneity were also present in mature cells.

Furthermore, the lineage effects were also manifested in cellular gene expression. Using the lineage-resolved scRNA-seq data, we found that transcriptome divergence between cells increased with cell lineage distance at both the 350-cell stage and the 600-cell stage, when most cells have completed embryonic cell differentiation (Figures S6B and S6C). In addition, using an expression atlas of 93 genes in 363 lineage-identity resolved cells (Liu et al., 2009), we confirmed the existence of the lineage effects on gene expression after hatching in L1 stage animals (Figure S6D).

Finally, since CAL changes with both lineage and tissue fates, we further determined the association between CAL and tissue fate after considering the influence of lineage. To control lineage distance, CAL divergences were compared between intra-tissue cells and the lineage distance-matched inter-tissue cells; this analysis showed that CAL divergences between intra-tissue cells were significantly lower than the control cells for all tissue types (Figure 5E and Table S5). Furthermore, the relative pair-wise CAL divergences between cells were quantified by normalizing CAL divergence to the average CAL divergence of all cells at the same lineage distance. Unsupervised clustering of cells using these lineage-effects controlled CAL divergence revealed five broad clusters, each of which was significantly enriched for cells of certain tissue types (Figures 5F and 5G, and Table S5). These results confirm a contribution of CAL convergence during tissue differentiation, even though the extent of convergence is affected by the lineage origin of cells.

In summary, while the regulatory states of cells with distinct lineage origins converge during tissue fate differentiation, the lineage history of cells is recorded in the cellular chromatin and gene expression programs that contribute to intra-tissue heterogeneity. This lineage-dependent tissue heterogeneity raises the possibility that the lineage origin of cells may drive the functional diversification of cell types within a tissue.

### A Predetermination Chromatin Strategy for Programming Left-Right Morphological Symmetry

With a bilateral body plan, many *C. elegans* cells belonging to the same tissue are organized in a left-right (L-R) symmetric morphological pattern following morphogenesis. In addition to being the same cell type, the vast majority of the cells on the left side are indistinguishable from their right-side counterparts in terms of anatomy and function (Sulston et al., 1983). L-R cells are generally organized as pairs of symmetric lineage groups, in which all cells located on the left are derived from one progenitor cell and the corresponding cells located on the right are derived from a different progenitor cell (Figure 6A).

**Figure 6.**
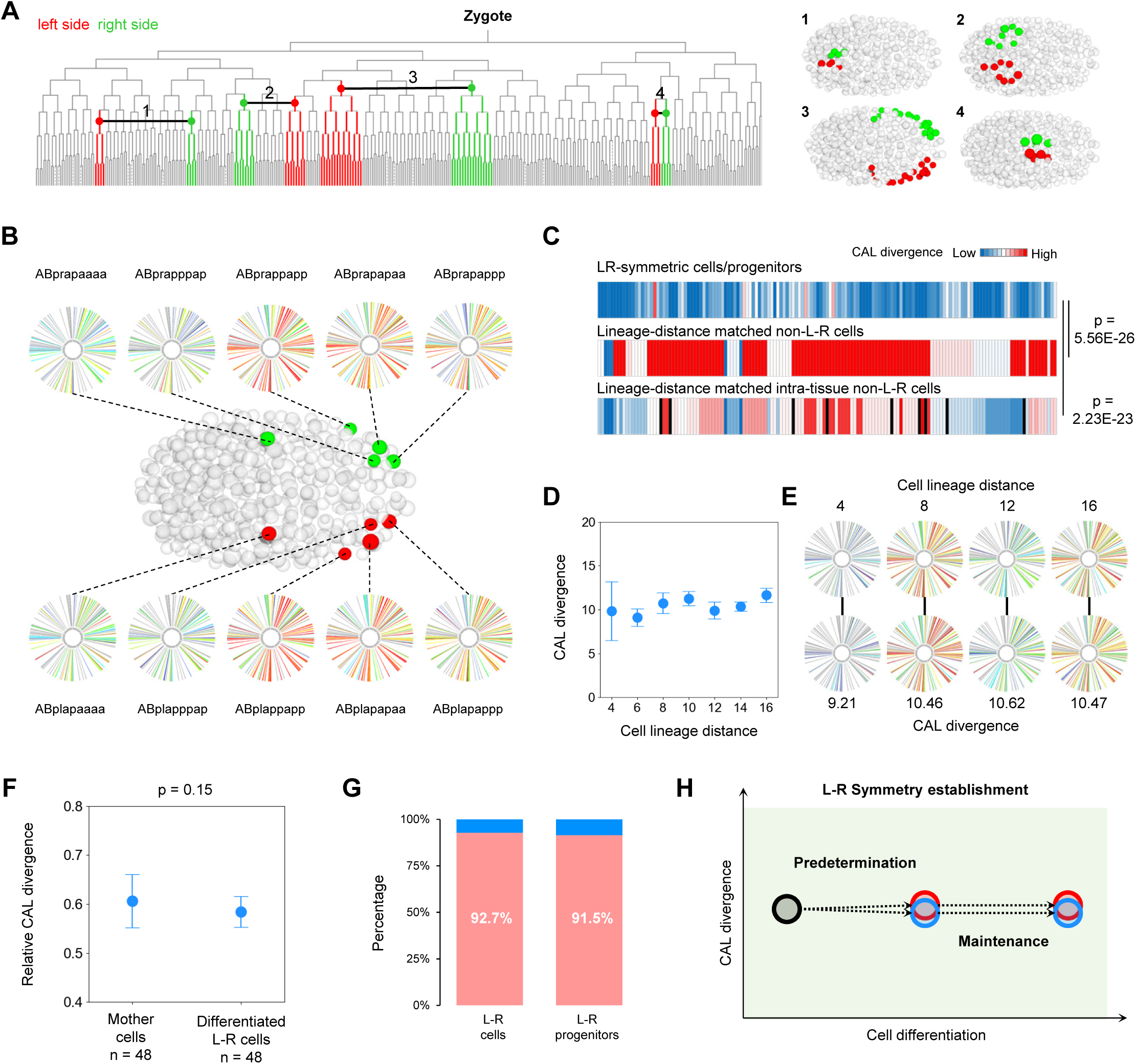
Predetermination of CAL during Establishment of L-R Symmetry. (A) L-R symmetry. Left: Four pairs of L-R symmetric cell lineages are linked by horizontal lines in the cell lineage tree. Right: embryonic locations of cells. (B) CAL of five representative pairs of L-R symmetric cell/progenitor cells. (C) Heatmap visualization of CAL divergences between cells in all pairs of L-R symmetric cells/progenitors and between control cells. Statistics: Wilcoxon signed-rank test, n = 149. (D) CAL divergences between L-R symmetric cells with different cell lineage distances. Data are presented as mean ± 95% CI. Statistics: Mood’s median test. (E) CAL and CAL divergences of representative pairs of L-R symmetric cells/progenitors with different cell lineage distances. (F) CAL divergences between differentiated L-R symmetric cells and between their mother cells. Data are presented as mean ± 95% CI. Statistics: Mann-Whitney U test. (G) Percentage of cases in which the cellular transcriptome is indistinguishable between differentiated L-R cells and between their progenitor cells. (H) The predetermination chromatin strategy to drive L-R symmetry establishment. See the main text. See also Figure S7 and Table S6.

Given the high functional similarity between L-R cells, we determined whether they exhibit higher CAL similarity than non-L-R cells. Some of the traced terminal cells are L-R cells or progenitor cells in the corresponding L-R symmetric lineages (L-R progenitors) (Figure 6B and Table S6). CAL divergence between the L-R cells/progenitors in each pair was significantly lower than the CAL divergence between the lineage-distance-matched non-L-R cells (p = 5.56E-26, Figure 6C). Furthermore, in 90% of cases, CAL divergence between L-R cells/progenitors was lower than the CAL divergence between the lineage-distance-matched intra-tissue cells (p = 2.23E-23, Figure 6C). Nevertheless, the L-R cells *per se* do not fully account for the higher CAL similarity observed between cells of the same tissue. CAL divergence between intra-tissue cells was significantly lower than that between inter-tissue cells after removing the L-R cells (Figure S7A). This pattern was also evident in the cellular gene expression data of L1 stage animals, in which gene expression divergences between L-R cells were significantly lower than those between other cells with a matched lineage distance within the tissue (Figure S7B and Table S6).

As in other intra-tissue cells, cells in many L-R pairs have distinct lineage origins. Surprisingly, distinct from other intra-tissue cells, we found that the lineage-dependent differences in CAL diminished between L-R cells (Figure 6D). As exemplified by Figure 6E, CAL divergence between L-R cells are independent of cell lineage distance, as the divergence between L-R cells/progenitors separated by eight generations (lineage distance = 16) are highly comparable to those separated by a less number of generations (lineage distance = 12, 8, and 4). Moreover, analysis of cellular gene expression in the L1 stage animals generated a similar pattern (Figure S7C). These results indicate that chromatin regulation of establishing L-R symmetry is distinct from that of general tissue differentiation.

Two possible mechanisms may underlie the difference. First, a similar CAL is predetermined in the progenitors of the corresponding L-R cells. Second, convergence of CAL is more robust during the establishment of L-R symmetry as compared to during general tissue differentiation. If the predetermination model is correct, we expected that CAL divergence between differentiated L-R cells would be similar to that between their earlier progenitor cells. From the traced cells, we identified 48 pairs of L-R cells that had completed embryonic mitosis, which represents cases where tissue differentiation and symmetry establishment have been completed. The degree of CAL divergence between these differentiated L-R cells and the degree of CAL divergence between their mother cells was highly comparable, which is consistent with the predetermination model (Figure 6F and Table S6). Furthermore, the analysis of cellular transcriptomes also supported the predetermination model. According to the cell identity assignment results of the scRNA-seq data (Packer et al., 2019), the transcriptomes of differentiated L-R cell pairs (92.7%) and L-R progenitor cell pairs (91.5%) were indistinguishable in the vast majority of cases (Figure 6G). These results favor predetermination of the cellular regulatory state during establishment of L-R symmetry.

In summary, the results of cellular CAL combined with transcriptome data reveal distinct chromatin strategies that are used to regulate the establishment of L-R symmetry as compared to general tissue differentiation (Figures 6H and 4G). During establishment of bilateral symmetry, a highly similar CAL is predetermined in corresponding early progenitors of prospective L-R symmetric cells and maintained during later development.

### Co-Regulation of Chromatin Activity Drives the Functional Coordination of the Genome

The cell-centric analyses described above revealed a multidimensional regulation of cell differentiation by chromatin. We sought to investigate the genomic organization of chromatin activity. Specifically, we asked what genomic regions that exhibit concordant inter-cell dynamics of chromatin activity (CDCA) predict the structural and functional relevance of the genome. We calculated the divergence of inter-cell dynamics of chromatin activity between 113 genomic positions and identified nine clusters (720 pairs) of CDCA regions (Figure 7A and Table S7). Some CDCA regions were linked on the same chromosome, at a distance of less than 1 Mb (Figure 7B). It is known that stretches of the genome are organized as a topology associated domain (TAD) in which chromatin interacts more frequently and usually exhibits similar properties and activities (Soler-Oliva et al., 2017). Indeed, using a previously generated dataset (Crane et al., 2015), we found that these pairs of CDCA regions were significantly enriched (2.65 fold) for those regions that are located within the same TAD (Figures 7C and 7D). This suggests that CDCA regions inform structurally related genomic regions.

**Figure 7.**
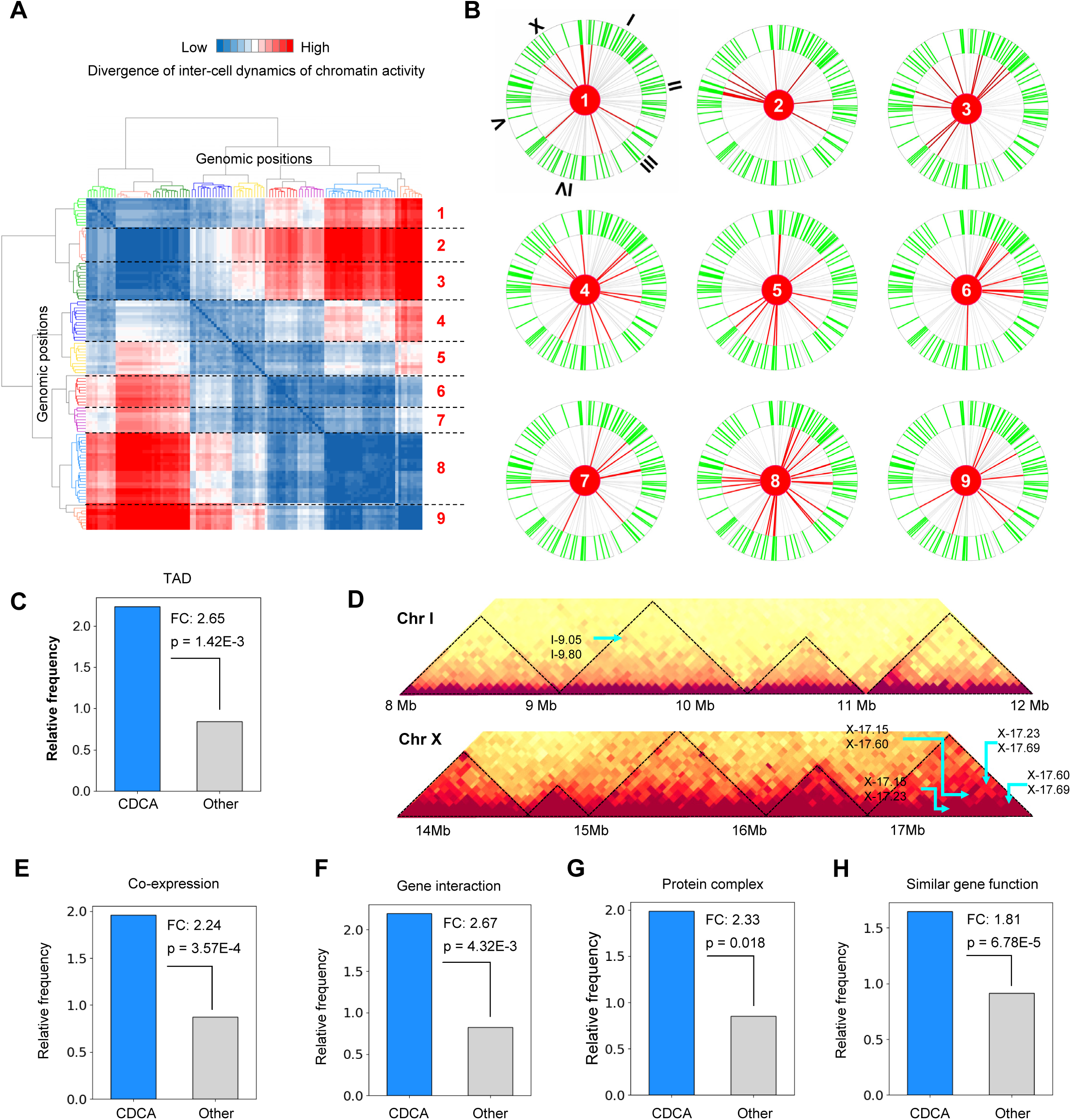
Concordant Inter-Cell Dynamics of Chromatin Activity Inform Structural and Functional Relevance of Genome Regions. (A) Hierarchical clustering of GFP integration sites based on the divergence of the inter-cell dynamics of chromatin activity. (B) Circos plots showing the genomic localization of nine clusters of CDCA regions. (C) Relative frequency of CDCA regions (blue) and non-CDCA regions (gray) in the TAD and non-TAD. (D) Examples showing the enrichment of CDCA regions in the same TAD. The heatmaps show the chromatin interaction maps binned at the 50-kb window for part of chromosome I and X. Triangles indicate the TAD regions identified in the original study and arrows indicate regions in the same CDCA clusters are located within the same TAD. (D-G) Relative frequency of co-expressed genes (D), interacted genes (E), genes present in the same protein complex (F), and genes with similar functions (G) in all genes located near the CDCA regions (blue, n = 720) and non-CDCA regions (gray, n = 5608). Statistics: Hypergeometric test. See also Table S7.

The vast majority of CDCA regions, however, are located far from each other or are even located on different chromosomes (Figure 7B), which raises the question as to whether they are predictive of co-regulated or functionally related regions? We tested this possibility by examining whether genes near the CDCA regions tend to be functionally related. Three types of functional relevance were tested: co-expression, gene interaction, and functional similarity, and all of the results supported the hypothesis (Figure 7E-7H and Table S7). First, pairs of CDCA regions were significantly enriched for genes that are co-expressed across greater than 900 conditions (2.24-fold enrichment, Figure 7E). Second, genes located near pairs of CDCA regions tended to interact with each other genetically, physically, or regulatorily (2.67-fold enrichment; Figure 7F) and were enriched for proteins that are present in the same complexes (2.33-fold enrichment; Figure 7G). Finally, genes near pairs of CCAD regions tend to encode proteins with identical functional annotations (1.81-fold enrichment; Figure 7H).

These genome-centric analyses reveal that co-regulation of chromatin activity across cells is widespread in the genome and it predicts topologically and functionally related genomic regions. This raises the possibility that chromatin regulation drives the functional coordination of genomic regions.

## DISCUSSION

### Single-cell Analysis of Positional Effects on Reporter Expression Reveals the Functional State of Chromatin During *In Vivo* Development

Many sequencing-based approaches efficiently dissect different aspects of the state of chromatin, such as DNA and histone modifications (Mikkelsen et al., 2007; Xie et al., 2013; Zhu et al., 2018), binding of regulatory proteins (Gerstein et al., 2010), accessibility (Cusanovich et al., 2018a), and spatial organization (Lieberman-Aiden et al., 2009). However, it is challenging to use these approaches to elucidate the functional state of chromatin in explicit single cells during *in vivo* development. First, the contribution of chromatin’s biochemical or biophysical states to its activity is highly multifaceted. For example, the influence of histone modification on gene expression is highly context-dependent and often relies on other types of histone modifications (Karlic et al., 2010; Wang et al., 2008). Similarly, high chromatin accessibility does not always correspond to high activity (Arnold et al., 2013). This uncertainty complicates the interpretation of the regulatory role of chromatin if only a limited number of chromatin properties are evaluated. Second, although the performance of some of the methods mentioned above has been improved for analyzing low cell numbers or the chromatin state in single cells (Ai et al., 2019; Cusanovich et al., 2018a; Cusanovich et al., 2018b; Stevens et al., 2017; Zhu et al., 2018), significant challenges remain with assigning lineage identities to the measured chromatin states in individual cells.

In this work, we quantified the positional effects on a reporter gene that is highly responsive to chromatin environments as a direct functional measurement of chromatin activity. Since the same gene was introduced into many genomic positions and ultimate gene expression was used as a readout of the chromatin state, the data described here represent the functional state of chromatin. Multiple pieces of evidence demonstrate that positional effects on gene expression reliably indicate chromatin activity (Figure 1). More importantly, systematic cell lineage tracing and single-cell quantification of gene expression enabled us to synthesize the CAL in lineage-resolved cells, which facilitated a precise analysis of the developmental role of chromatin. Due to low-throughput generation of randomly integrated transgenic animals, the genomic resolution of the present CAL is not high (0.88 Mb). In the future, this limitation will be improved by using CRISPR/Cas9-mediated genome editing to integrate reporter genes into a specific region of interest or throughout the genome at a higher resolution. In addition, the expression window of the *eef-1A.1* promoter in the integrated strains does not cover early embryonic cells, which prevented us from measuring cellular chromatin during very early embryogenesis. This limitation can be resolved by using other promoters that are responsive to chromatin environments and expressed ubiquitously during very early development. Nevertheless, the functional nature, cellular resolution, and high cell coverage of the CAL described here provides a unique opportunity to systematically explore the role of chromatin in development at single-cell resolution. With suitable promoters, chromatin activity at any genomic position in any cell and at any developmental stage will be traceable using our single-cell approach.

### Chromatin State Encodes Regulatory Information of Cell Lineage Differentiation

While chromatin state provides rich information to regulate gene expression and to specify cellular regulatory states, to what extent the regulatory processes that govern *in vivo* development are encoded in the chromatin of individual developing cells remains an open question. Lineage-resolved CAL combined with the extensive prior knowledge of single-cell biology of *C. elegans* development allows us to investigate chromatin regulation of cell lineage differentiation in single cells. We analyzed the dynamics of CAL across three critical dimensions of cell differentiation regulation: cell lineages, tissue fate, and symmetric morphological organization (Labouesse and Mango, 1999). These single-cell analyses are used to evaluate the general properties of chromatin dynamics during cell differentiation. At the lineage level, CAL changes steadily in general and rapidly following specific cell divisions, which is predictive of the kinetics of lineage-coupled cell differentiation and anterior-posterior fate asymmetry. Consequently, characteristic CAL is established in cells according to their lineage origin (Figures 2 and 3). At the tissue level, similar CALs are specified in cells that are from multiple lineages when they differentiate into the same tissue type, especially for the cells that are organized in an L-R morphologically symmetric pattern (Figures 4-6). Accordingly, the characteristic CAL is specified to cells according to the tissue type. The dynamic patterns further demonstrate that many critical regulatory processes of cell differentiation are readily inferable from cellular chromatin, including lineage specifications, anterior-posterior fate asymmetry, tissue fate specification, cell heterogeneity, and bilateral symmetry establishment. A previous study that profiled genome-wide maps of DNase I-hypersensitive sites in diverse human embryonic stem cells and adult primary cells revealed that the chromatin landscape defines cell lineage relationships, cell fates, and cellular maturity (Stergachis et al., 2013). Our findings combined with this previous study (Stergachis et al., 2013) suggest that cellular chromatin states encode informative regulatory signals that drive *in vivo* cell lineage differentiation.

### Distinct Chromatin Strategies Regulate Tissue Fate Differentiation and Establishment of L-R Symmetry

We found that different chromatin strategies are invoked in order to specify similar regulatory states in cells of the same tissue type. In general, a convergence strategy is used to progressively specify similar CALs and gene expression programs in cells from multiple lineages according to the future tissue types that they differentiate into. However, for those that are organized in an L-R morphologically symmetric pattern, a predetermination strategy is used to establish highly similar CALs and gene expression early in the corresponding precursors of the left and right cells.

While the convergence strategy specifies similar CALs in cells according to tissue type, intra-tissue cells exhibit considerable CAL and gene expression heterogeneity, which is lineage-origin-dependent. This suggests that differences in developmental histories of cells contribute to intra-tissue/cell-type heterogeneity. A similar finding was observed in adult mouse tissue, in which the same cell type exhibited characteristic chromatin accessibility profiles according to its location in the body (Cusanovich et al., 2018a). While it remains to be determined, these cells are likely to be from discrete cell lineages. Thus, these findings support the existence of a widespread lineage-dependent functional organization of cells at the sub-tissue/cell-type level. Consequently, intra-tissue cells with a closer lineage relationship are more functionally related and exhibit lower heterogeneity in the regulatory states. Consistently, as shown by previous studies, lineage related neurons are more functionally related in terms of microcircuit assembly and stimulus feature selectivity (Li et al., 2012; Yu et al., 2012). The progressive convergence of CAL during tissue differentiation also implies that the pioneer transcription factors responsible for specifying tissue fate may play an essential role in shaping the characteristic chromatin landscape, and that the cooperative interactions between chromatin and pioneer factors drive tissue differentiation (Zaret and Mango, 2016). Consistent with these conclusions, loss of function of *elt-1*, a specifier of skin fate, induced significant changes in chromatin activity (Figure 4E). Important follow-up studies include: identifying which pioneer factors are responsible for remodeling the chromatin landscape; how these pioneer factors co-specify chromatin state in functionally related cells from discrete lineages; and determining how the lineage-dependent heterogeneity in chromatin is established.

Conversely, a predetermination strategy is used to establish a highly equivalent CAL between the cells that are organized in an L-R symmetric pattern. We propose that the predetermination strategy ensures that highly equivalent regulatory states are robustly specified to prospective L-R cells so that identical cellular functions are performed on different sides of the body. A small number of morphologically L-R symmetric cells exhibit functional asymmetry (Hobert, 2014). Interestingly, a previous study that focused on a pair of functional asymmetric neurons (ASEL/ASER) reported that, although functional asymmetry arises very late during embryogenesis, differential regulatory states are predetermined at a very early stage in the corresponding precursor cells through differential chromatin decompaction of the *lsy-6* miRNA locus (Cochella and Hobert, 2012). It appears that predetermination of the chromatin state is a widely adopted strategy used to regulate functional symmetry and asymmetry between L-R cells. An intriguing question that should be addressed is understanding when and how equivalent CALs are precisely specified in early progenitor cells that are not lineage-related. Since cells from different cell lineages generally exhibit divergent CALs, the specification of equivalent CALs would be an actively regulated process. In addition, considering that progenitors of L-R symmetric cells originate from multiple cell lineages and occupy different regions of the embryo, the lineage-based cell-autonomous mechanisms may play a leading role in this process.

## ACKNOWLEDGMENTS

This work was supported by grants from the “Strategic Priority Research Program” of the Chinese Academy of Sciences to Z.D. (XDB19000000), the National Natural Science Foundation of China to Z.D. (31722035 and 3177090082), and the State Key Laboratory of Molecular Developmental Biology, China. Some strains were provided by the CGC, which is funded by the National Institutes of Health Office of Research Infrastructure Programs (P40 OD010440).

## AUTHOR CONTRIBUTIONS

Z.D., Z.Z. and R.F. conceived the project and designed the study; R.F., W.X., Y.W. and X.M. conducted the experiments and generated the data; Z.Z. and Z.D. performed the data analysis and statistics; Z.D. wrote the manuscript with input from all authors.

## DECLARATION OF INTERESTS

The authors declare no competing interests.

## Supplemental Information

### 1. CONTACT FOR REAGENT AND RESOURCE SHARING

Further information and requests for resources and reagents should be directed to and will be fulfilled by the Lead Contact, Zhuo Du

### 2. EXPERIMENTAL MODEL AND SUBJECT DETAILS

#### 2.1 *C. elegans* Strains and Culture

Genotypes of all *C. elegans* strains used in this study are listed in Table S1. Some of the strains were obtained from the Caenorhabditis Genetics Center. Unless otherwise specified, all strains were cultured in incubators at 21 °C on nematode growth media plates seeded with OP50 bacteria.

### 3. METHOD DETAILS

#### 3.1 Selection and Verification of Reporter Strains

From a previously generated collection of integration strains (Frokjaer-Jensen et al., 2014) (http://www.wormbuilder.org/reagents/fluorescent-marker-strains/), a total of 116 report strains were selected based on the following criteria: (i) driven by the *eef-1A.1* promoter (previously known as *eft-3*), (ii) use of GFP as the fluorophore, (iii) containing a nuclear localization signal, and (iv) use of a unique integration site. Three strains (EG8880, EG8912, EG8860) exhibited severe growth and developmental defects and were removed. Each of the 113 strains was crossed with the JIM113 strain that ubiquitously expressed mCherry in the nucleus for lineage tracing, which resulted in a collection of 113 dual-fluorescent reporter strains (Table S1).

The integration sites of P*eef-1A.1*::GFP in all transgenic strains had been characterized previously. To further ensure the accuracy of the integration sites and copy number, 18 strains were randomly selected and the integration sites were reexamined using inverse PCR following a previously established protocol (Frokjaer-Jensen et al., 2014). Genomic DNA was extracted and digested overnight with the restriction enzyme DpnII that cut at a unique site located in the MOS 1 vector and potential sites located near the integration site in the *C. elegans* genome. Next, the digested DNA was circularized by T4 ligase during a 2-4 hour incubation. Two rounds of PCR were performed with the circularized DNA containing flanking sequence as the template and with two pairs of nested primers targeting the MOS 1 sequence. PCR products were gel-purified and sequenced. The sequences were then aligned to the *C. elegans* genome (WBcel235/ce11) in order to identify the sequence of the regions flanking the integration site. For all 18 tested strains, integration sites were identical or within 1 kb of the previous annotation; additionally, only one integration site was identified in all examined strains (Figure S1A).

#### 3.2 Embryo Mounting and 3D Time-Lapse Imaging of Embryogenesis

##### Embryo Mounting

A previously established procedure was used to prepare and mount early *C. elegans* embryos, with minor modifications (Bao and Murray, 2011). Briefly, six young adult worms with one row of eggs in the gonad were picked and transferred onto a Multitest slide (MP Biomedicals) with a droplet of M9 buffer, then the worms were cut open to release early embryos. Two- to four-cell stage embryos were identified under a dissecting microscope (Nikon SMZ745) that were then transferred into a droplet of egg buffer (∼2 µL) containing 20 μm polystyrene microspheres (Polyscience) on a coverslip (Fisherbrand) using an aspirator tube assembly (Sigma-Aldrich). The various positions of embryos were adjusted so that they were arranged in a few clusters, with each cluster containing two to three embryos (fit with one imaging field). Finally, an 18 × 18 mm coverslip was placed on top of the droplet and the slide was sealed with melted Vaseline.

##### Live Imaging

3D time-lapse imaging was performed using a spinning disk confocal microscope (Revolution XD) with an inverted microscope body (Olympus IX73), a spinning-disk unit (Yokogawa CSU-X1,) XYZ stage with Piezo-Z positioning (ASI PZ-2150-XYZ), an integrated solid-state laser engine (Coherent; 50 mw at 488 nm and 50 mw at 561 nm), and an Electron Multiplying Charge-Coupled Device (EMCCD; Andor iXon Ultra 897). Images were taken at 20 °C using the multidimensional acquisition module of MetaMorph software (Molecular Devices) under a 60X objective (PLAPON 60XO, N.A. = 1.42). Images were recorded for 240 time-points at a temporal resolution of 75 seconds, and at each time point, three slide positions with two to three embryos were scanned for 30 Z focal planes with 1 µm spacing. Laser power and exposure time for mCherry (561 nm) and GFP (488 nm) were optimized in order to minimize photo-damage while maintaining a high signal to noise ratio. Laser power for both mCherry and GFP was increased 3% for every Z plane when the focal plane went deeper into the samples in order to partially compensate for the decay of the fluorescence signal over Z focal panes. Laser power and exposure time used for mCherry and GFP for the first Z plane were 0.08 for 50 milliseconds and 0.08 for 20 milliseconds, respectively. All wild-type embryos (n > 50) imaged with this parameter hatched at a time that was highly comparable to embryos without laser excitation without obvious phenotypic abnormality. For individual time points, the two-channel image series were organized as 3D tiff stack images and were directly used for cell identification, tracing, lineage construction, and quantification of single-cell reporter expression.

#### 3.3 RNAi

RNAi experiments were performed using a standard feeding protocol (Kamath et al., 2001). All RNAi clones were from the *C. elegans* RNAi feeding library constructed by Julie Ahringer’s group (Source BioScience). The insert of individual clones used in this study was verified by end sequencing. For RNAi treatment, ten worms synchronized at the L1 stage (P0) were transferred to RNAi plates containing 3 mM IPTG and that had been seeded with the corresponding bacteria expressing the double-stranded RNA against the target gene. After several days of RNAi exposure, embryos of P0 or P1 animals were subjected to imaging and analysis.

#### 3.4 Lineage Fate Transformation and Tissue Fate Perturbation Experiments

##### Induction of Lineage Fate Transformation

Two pairs of cell lineages were selected, ABalp-ABarp and MS-E, and multiple reporter integration strains exhibiting differential GFP expression in descendant cells from corresponding lineages were used to examine the relationship between CAL and lineage fate. Transformation of the ABalp lineage fate to that of normal ABarp was induced by RNAi targeted against the *lag-1/CLS* gene. It was previously established that the fates of ABalp and ABarp lineages are distinguished by Notch signaling, and a perturbation of Notch singling components induces ABalp-to-ABarp lineage fate transformation (Du et al., 2014; Moskowitz and Rothman, 1996). Transformation of MS lineage fate to that of normal E was induced by RNAi targeted against the *pop-1/TCF* gene. The distinguishing of fate between MS and E lineages is regulated by Wnt signaling. Loss of function of the Wnt effector *pop-1/TCF* transforms the developmental fate of MS lineage to that of the E lineage (Du et al., 2014; Lin et al., 1995). RNAi treatments were performed on seven (*lag-1* knockdown) and five (*pop-1* knockdown) strains in which GFP was integrated into distinct genomic positions. Only embryos showing evidence of lineage fate transformation were used for comparing GFP expression between cells from corresponding lineages.

##### Perturbation of Tissue Fate

Perturbation of skin fate was performed by using a loss of function mutant of *elt-1/GATA1,* a skin fate specifier. As shown previously, the conserved GATA transcription factor *elt-1* is necessary and sufficient for specifying skin cell fate. Loss of *elt-1* leads to extra neuronal (from the ABarp lineage), muscle (from Caa and Cpa lineages), and other cell types (from the ABpla lineage) at the expense of skin cells (Page et al., 1997). Furthermore, forced ectopic expression of *elt-1* is sufficient to transform most embryonic cells into skin cells (Gilleard and McGhee, 2001). We used the *elt-1(zu180)* allele, which contains a stop codon within the first zinc finger domain, to induce fate changes of skin cells. To examine the influence of tissue fate perturbation on CAL, we compared changes in GFP expression in cells from three major skin lineages (ABarp, ABpla, and ABpra) using three integration strains in which GFP is expressed in cells from the aforementioned lineages.

### 4. QUANTIFICATION AND STATISTICAL ANALYSIS

#### 4.1 Cell Identification, Lineage Tracing, and Manual Curation

Image series were processed with StarryNite software (Santella et al., 2014; Santella et al., 2010) in order to reconstruct de novo the embryonic cell lineage by automated cell identification and tracking. Automated cell identification was performed using a hybrid blob-detection algorithm in order to segment individual nuclei in the 3D image stacks based on the ubiquitously expressed mCherry fluorescence signal that localizes to the nucleus (Santella et al., 2010). Next, automated cell tracing was performed using a semi-local neighborhood-based framework to link all cells at a preceding time point to those at the subsequent time point, and if cell division occurs, a mother cell is linked to two daughter cells (Santella et al., 2014).

Second, raw cell identification and tracing results were manually inspected and curated systematically to ensure high accuracy using the AceTree software (Katzman et al., 2018). While the accuracy of automated cell detection and tracing of StarryNite software is high (> 99%), the accumulative nature of the errors affect the accuracy of cell lineage results (Santella et al., 2014). Hence, a systematic correction of lineage errors, especially those that occur in the early developmental stage, is indispensable. AceTree software provides an interface for visualizing the traced cell lineage as a binary tree structure and links all cells on the tree to the raw 3D images. With this function, users can identify potential lineaging errors in the tree, inspect the cell relationships on the raw images, and finally modify the identification or tracing results when necessary. Because the *C. elegans* cell lineage is invariant, the vast majority of errors can be efficiently captured by visual or computational screening of the unusual lineage topologies. Detailed procedures for error detection and correction were described previously (Du et al., 2015).

Third, the lineage identities of all traced cells were determined and assigned a unique name according to Sulston’s nomenclature (Sulston et al., 1983). Cell identities were first determined for all cells (ABa, ABp, EMS, and P_2_) in the 4-cell stage embryos based on the stereotypical arrangement of cells in the embryo and the timing of cell divisions. Specifically, ABa and P_2_ are located in the anterior and posterior parts of the embryos, respectively, and ABa and ABp divide earlier than EMS and P_2_ cells. Then, the names of their descendants were determined according to the mother cell name and cell division pattern. All cell divisions fall into three broad categories: anterior-posterior (a/p), left-right (l/r), and dorsal-ventral (d/v), according to the orientation of cell division relative to the body axis. In general, the full name of a mother cell is propagated to the daughter cells with an additional letter specifying the cell position of the daughter cell relative to the body axis following cell division of the mother. For example, ABal specifies the daughter cell of ABa that is located on the left side following the l/r division of the ABa cell, and ABala specifies the daughter cell of ABal that is located anteriorly following the a/p division of the ABal cell. Except for the few early progenitor cells (MS, E, C, D, P_3_, P_4_, Z2, and Z3) to which a particular name was assigned to highlight their developmental properties, the name assignment of all cells followed this general rule. Detailed nomenclature information is described elsewhere (Santella et al., 2010; Sulston et al., 1983). Using the aforementioned image bioinformatics, the embryogenesis was digitized at cellular resolution and the cell lineage identities of all traced cells were determined.

#### 4.2 Quantification of Reporter Expression in Lineaged Cells

##### Quantification of raw GFP expression

Each strain that was used to quantify the positional effects of reporter expression carried two nucleus-localized fluorescent proteins. The ubiquitous mCherry was used for the aforementioned cell lineage reconstruction, and the fluorescent intensity of GFP integrated into a specific genomic position was used to measure the reporter expression level in each traced nucleus simultaneously. The segmentation of each nucleus at each time point and tracing of nuclei across time during the lineage construction step facilitated a direct quantification of GFP expression in each lineage-resolved cell.

First, we measured raw GFP expression in each cell. GFP expression was calculated as the average intensity of all pixels within each identified nucleus at each time point minus the average intensity of local background. The average pixel intensity in the center Z plane of each nucleus was used to approximate the expression level in the nucleus. The background signal for each nucleus was estimated using a previously described method in which the average pixel intensity was calculated within an annular area between 1.2-radius to 2-radius from the centroid of the nucleus. Nearby nuclei that overlapped with the annular area were not included in the background measurement (Murray et al., 2008). GFP intensities of the same cell at multiple time points were averaged that represented cellular average GFP expression abundance. Because the nuclei morphology at the time point before and after a cell division is not spherical (which would affect the accuracy of intensity measurements), the values at these time points were excluded when calculating the average GFP expression in a cell.

##### Compensation for Depth-dependent Attenuation of Fluorescence Intensity

GFP levels were adjusted by compensating the depth-dependent attenuation of fluorescence intensity. A fundamental problem associated with 3D fluorescence confocal imaging is the attenuation of light with depth, which is caused by the absorption and scattering of both the excitation and fluorescence light (Kervrann et al., 2004). Consequently, without adjustment, the measured fluorescence intensities of cells reside deeper in the embryo (far from the microscope objective) and are significantly weaker than those located in shallower slices. This effect could significantly obscure a reliable comparison of GFP expression between single cells in an embryo, since the equivalent cell are located at different depths (Z planes) relative to the objective during imaging (Figure S1B).

Although the attenuation effect was partially compensated for during imaging by increasing the laser power of the excitation light with depth, this effect was still present in the acquired images. This phenomenon is best illustrated by comparing GFP expression in equivalent cells between embryos of the same strain (experimental replicates) that are oriented differently during imaging. There are two types of embryo orientations at the 350-cell stage in which either the ventral or the dorsal side is placed near the objective (termed VNO and DNO, respectively). As shown in Figure S1C, cellular GFP intensity was highly consistent between embryos with identical orientation (average Pearson correlation coefficient r = 0.85). However, the intensity differs considerably between embryos with different orientations (average Pearson correlation coefficient r = 0.22), especially for those cells located far from the center Z planes of the embryo.

This discrepancy allowed us to model and correct the residual attenuation effect. Using the information of each cell’s Z position and GFP intensity, we applied various attenuation factors per Z plane (α) in order to adjust the GFP intensity at any Z plane to the center plane (Z = 15) using the equation I_i_ = I_c_ · (1 + α)^(i-c)^, where I_i_ and I_c_ specify GFP intensity of the cell at plane i and the center plane, respectively. The performance of each α was evaluated by quantifying the correlation coefficient of cellular GFP expression between experiment replicates with different embryonic orientations. We used 49 embryos of 13 transgenic strains to model the performance of adjustment and determined whether the attenuation effect depended on the magnitude of fluorescent intensity. We found that the attenuation was significant when the cellular GFP intensity was greater than seven and that α = 0.054 yielded the best performance regarding adjustments of the attenuation (Figure S1D). These parameters were applied to all cells in all embryos, which dramatically removed the residual attenuation effects (see Figure S1E for representative examples). This adjustment of GFP intensity facilitated a reliable inter-cell comparison of GFP expression in an embryo, especially when the cells are located in significantly different Z planes.

##### Quantification of Instantaneous GFP Expression

We determined instantaneously expressed GFP in each cell. Due to the high stability of GFP protein (with a half-life of over 20 hours in mammalian cells) and a fast cell cycle progression during embryogenesis (median cell cycle length = 42 minutes until the 350-cell stage), the measured cellular GFP intensity consisted of both GFP inherited from the previous cell cycle (expressed in the mother cell) and GFP expressed in the present cell. Since GFP expression was continuously imaged at a high temporal resolution, the inherited and newly expressed GFP was distinguished by subtracting the GFP intensity of mother cells from the corresponding daughter cells. The GFP intensity value at the time point before the cell division of the mother cell represented the GFP expressed in the mother cell and was subtracted from the value of the daughter cells, which assumed no gene expression occurred during cell division.

##### Determination of Expression Cut-off

The cut-off for calling GFP expression was determined. Only strains with the mCherry lineaging marker but not the GFP transgene were used as the control and cellular GFP intensity was quantified in 20 embryos to model the distribution of GFP intensity in non-expression cells. A cut-off of 6.36 (q < 0.01) at which the false discovery rate was 6.9E-5 for cells in control embryos was used to refine GFP expression levels. For cells with an intensity lower than the cut-off, the expression level was set to zero; otherwise, the cut-off value was subtracted from the cellular GFP intensity. These values were then log_2_(X+1) transformed to represent the final cellular instantaneous expression level (Table S2).

#### 4.3 Synthesis of Single-Cell CAL

Taking advantage of the invariant cell lineage, we constructed CAL of individual cells by integrating the expression levels of the same reporter integrated into 113 genomic positions in cells with the same lineage identity (equivalent cells). Because cellular GFP expression for each integration line was quantified for multiple embryos (ranging from 2 to 8), to integrate the expression levels, only cells in which GFP is expressed in more than 60% of the embryo were considered to be expressing cells, and the values were averaged to represent consensus chromatin activity at a given genomic location in a cell (Table S2).

#### 4.4 Association of Position-Effects on GFP Expression with Chromatin Features

We systematically examined the biological relevance of the measured positional effects on GFP expression to chromatin activity and confirmed that GFP expression is highly consistent with many features of chromatin that are related to its activity (Table S2).

First, the positional distribution of GFP expression levels was consistent with the known genomic organization of chromatin activity. Similar to previous findings (Frokjaer-Jensen et al., 2016), GFP expression levels were significantly higher when integrated into autosomes as compared to the X chromosome (Figure S2B), and were significantly higher when integrated into the central regions of chromosomes as compared to the arm regions (Figure S2C).

Second, positional effects on GFP expression were consistent with the biochemical properties of chromatin that are related to its activity. Specifically, the correlation between GFP expression and histone modification, a key determinant of chromatin activity, was examined. We predicted GFP expression levels at each integration site using the combination of 19 types of histone modifications (Ho et al., 2014). Histone modification datasets were downloaded from the ModENCODE project website, and only histone modifications with embryonic datasets (n = 56) were used to ensure a comparison of histone modification and GFP expression at the comparable developmental stage. Each dataset was normalized by calculating the Z score Z = (x - µ)/α for each region across the genome, where µ denotes the mean and α denotes the standard deviation of the modification levels. The level of each histone modification at the transgene integration site was determined and was used to represent the modification level of each *Peef-1A.1::GFP* cassette by averaging the modification level of all genomic regions that overlapped with a 1-kb region centered on the integration site of the expression cassette. GFP expression levels were predicted by a linear model that considers all types of modifications (Table S2). If multiple datasets are available for the same histone modification, the dataset that yielded the best performance was used. The result showed that the values predicted by histone modifications correlated significantly with the measured values (Figure S2D, R = 0.82, p = 8.27e-29).

Third, position-effects on GFP expression were consistent with chromatin accessibility. Accessibility chromatin regions were defined based on a previous study in which ATAC-seq was applied to identify accessible chromatin at several developmental stages (Janes et al., 2018). An integration site was defined as a location in accessible chromatin if an ATAC-seq peak overlapped with a 1-kb region centered on the integration site (Table S2). We found that GFP expression was significantly higher when integrated into accessible chromatin regions as compared to the rest of the genome (Figure S2E).

Finally, position-effects on GFP expression were consistent with the 3D organization of chromatin that is known to affect its activity. Chromatin regions attached to the nuclear lamina tend to be heterochromatic and exhibit low transcriptional activity (Kind et al., 2013; Reddy et al., 2008). We examined whether GFP expression was reduced when located in the lamina-associated domains (LAD). LAD information was obtained from a previous study that determined regions associated with LEM-2, an inner nuclear membrane protein, in mixed-stage *C. elegans* embryos (Ikegami et al., 2010) (Table S2). Indeed, we found that when the transgene was located in the LAD lower expression levels were observed as compared to non-LAD regions (Figure S2F).

Collectively, comparison of GFP expression to the associated genomic organization and to the biochemical and biophysical properties of chromatin confirmed that positional effects on P*eef-1A.1*::GFP can serve as a reliable sensor of chromatin activity.

#### 4.5 Association of Position-Effects on GFP Expression with Genetic Features

##### Classification of Intragenic/Intergenic Regions

An integration site was defined as being located in the intragenic region if the annotated integration site is located within the gene body of any protein-coding gene; otherwise, the reporter was classified as being located in the intergenic region (Table S2).

##### Comparison of GFP Expression With Nearby Endogenous Genes

The endogenous gene expression dataset that was used was from a previous study that generated a whole-embryo time-course transcriptome at high temporal resolution (Hashimshony et al., 2015). Gene expression at stage 190 min past the first cell division that was comparable to the 350-cell stage was used to represent endogenous gene expression. For each transgene integration site, the expression levels of all endogenous genes whose transcription start site is located in an interval centered on the transgene integration site were averaged to represent the expression potential of nearby genes. The transcript abundances were log_2_(X+1) transformed, and intervals of various sizes ranging from 5 kb to 2 Mb were analyzed (Table S2).

#### 4.6 Analysis of the Dynamics and Implications of Cellular CAL in Cell Lineage Differentiation

##### Calculation of Cell lineage Distance

A previously described strategy was used to calculate the lineage distance between two cells as the number of cell divisions that separate these two cells from their lowest common ancestor (Du et al., 2015) (Figure S3A).

##### Definition and Quantitative Comparison of Developmental Fates Between Progenitor Cells

The developmental fate of a progenitor cell was defined retrospectively as the combinatorial pattern of tissue types it produced following development (Figure S3B). The 671 embryonic terminal cells were first classified into seven tissue/organ categories: neuronal system (Neu; including neurons and glial cells), pharynx (Pha), skin (Ski), body wall muscle (Mus), intestine (Int), and germ cell (Ger). Cells that did not fall into any of these categories and that undergo programmed cell death were not considered in the analysis. The developmental fate of each progenitor cell was expressed as a pattern describing the tissue type of each terminal cell in the lineal order. Similarity was quantified by aligning the two patterns and comparing tissue fates between each lineal equivalent terminal cell (e.g., comparison of Xapap to Yapap cells between progenitor cell X and Y). A score of 1 was assigned if two lineal equivalent cells belonged to the same tissue type; otherwise, a score of 0 was assigned. The similarity scores were averaged across all terminal cells and were used to represent overall fate similarity. In cases where two progenitor cells generated different numbers of terminal cells due to different rounds of cell divisions in intermediate cells, the number of cells and tissue fates in the smaller lineage was expanded accordingly in the corresponding branches to match the larger one. This strategy is illustrated in Figure S3B.

##### Quantification of CAL Divergence

CAL divergence between cells was quantified as the Euclidean distance between chromatin activity (GFP expression levels) across all integration sites using the equation 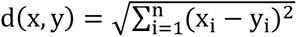, where x and y denote the vector of chromatin activity across 113 positions in the corresponding cells.

##### Identification of Transition Points of CAL

A retrospective approach was used to compare CAL divergence between cells from two daughter cell lineages to determine whether division of the mother cell leads to significant changes in CAL between two daughter cells. For each pair of daughter cell lineages generated by a progenitor cell, CAL divergences were calculated between cells from each daughter cell lineage (intra-daughter-lineage CAL divergences) and were compared to CAL divergences between cells from different daughter cell lineages (inter-daughter-lineage CAL divergences) (Figure S3E). A progenitor cell division was defined as a transition point of CAL if the inter-daughter-lineage divergences were significantly larger (Mann–Whitney U test, Benjamini-Hochberg adjusted p < 0.05) than any of the intra-daughter-lineage divergences (Table S3).

##### Clustering of Cells

Pair-wise CAL divergences (Euclidean distances) were first calculated between 364 traced terminal cells, and the CAL divergence between each cell pair of cells was then normalized to the mean CAL divergence of all cell pairs at the same lineage distance, which controlled for the influence of lineage distance between cells on CAL divergence. Groups of cells with similar CAL were identified by hierarchical clustering using the normalized cell-cell CAL divergence matrix and the following parameters: Euclidean distance as the distance metric, average linkage clustering as the linkage selection method, and distance threshold = 5.2 (Table S5).

##### Quantification of Gene Expression Divergence Between Cells

For single-cell transcriptome data (Packer et al., 2019), the Jenssen-Shannon distance was measured in order to quantify divergence, as was done in a previous study (Packer et al., 2019). Unless otherwise noted, only cells that had been assigned a unique or two possible lineage identities were used for analysis. For the identity-resolved single-cell gene expression dataset at the L1 stage (Liu et al., 2009), divergence was quantified by measuring Euclidean distance using the binary expression matrix.

##### L-R Symmetric Cells

The list of L-R symmetric cells and progenitor cells was defined by (Sulston et al., 1983) (Table S6).

#### 4.7 Analysis of the Genomic Organization of Chromatin Activity

##### Identification of Regions Exhibiting Concordant Inter-Cell Dynamics of Chromatin Activity (CDCA)

Pair-wise divergences (Euclidean distance) of chromatin activity among cells were calculated between all reporter integration sites. The matrix was used to identify clusters of genomic positions by hierarchical clustering using the following parameters: Euclidean distance as the distance metric, average linkage clustering as the linkage selection method, and distance threshold = 65 (Table S7).

##### Topologically Associating Domain (TAD) data

TAD information was obtained from a previous study that constructed a genome-wide chromatin interaction maps in mixed-stage *C. elegans* embryos (Crane et al., 2015).

##### Co-Expressed Genes

The comprehensive dataset of global gene expression changes across diverse genetic and environmental conditions (15,204 genes across 979 conditions) was from a previous study (Stuart et al., 2003). Pair-wise Spearman rank correlation of gene expression across all conditions was calculated between all genes, and a pair of genes was defined as being co-expressed if the correlation coefficient ranked among the top one-third of all values. Genes located within a 20-kb region centered on GFP integration sites were used to analyze the relative enrichment of co-expressed genes between CDCA regions (Table S7).

##### Gene interactions

Two sources of gene interaction data were used. First, a curated list of gene interactions (including genetic, physical, and regulatory interactions) was downloaded from WormBase (c_elegans.PRJNA13758.WS270.interactions.txt.gz). Second, a dataset of 612 putative *C. elegans* protein complexes generated by analyzing large-scale chromatographic fractionation-mass spectrometry (CF-MS) data was used to analyze physical interactions between genes (Hu et al., 2019). Due to a small number of gene interactions, genes located within a 50-kb region centered on the GFP integration sites were used to analyze the relative enrichment of gene interactions between CDCA regions (Table S7).

##### Genes with Similar Function

Functional annotations of all *C. elegans* genes were downloaded from the WormCat database, in which all *C. elegans* genes are classified into hundreds of non-redundant functional categories based on physiological functions, molecular functions, phenotypes, subcellular location, and other information (Holdorf et al., 2020). We used the Category 3 annotation (455 specific categories) and excluded four categories annotated as unknown for the analysis. Genes located within a 20-kb region centered on the GFP integration sites were used to assess the relative enrichment of genes with similar function between CDCA regions.

### 5. DATA AND SOFTWARE AVAILABILITY

All datasets supporting this article can be found in the supplementary data and online at http://dulab.genetics.ac.cn/sc-CAL. Live imaging data are available upon request.

### 6. SUPPLEMENTARY FIGURE LEGENDS

**Figure S1.**
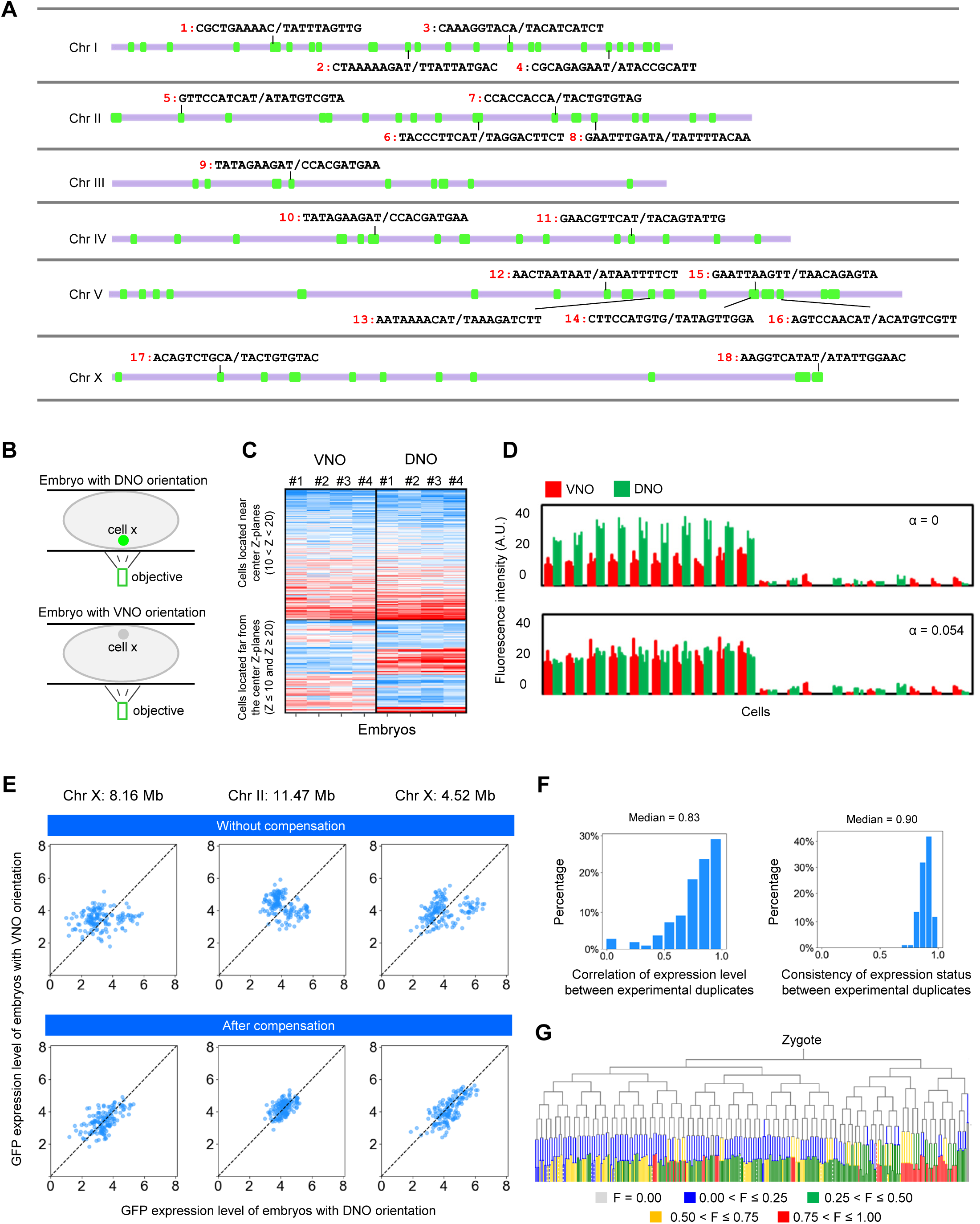
Quantification of Positional Effects on GFP Expression in Single Cells During *C. elegans* Embryogenesis, related to STAR Methods and Figure 1. (A) Verification of the integration sites of reporter lines. The identified genomic sequences flanking the integration sites are shown for each examined integration strain. (B) Representation of fluorescent intensity attenuation with depth. Due to different embryo orientations, the same cell in different embryos is located at a different depth relative to the microscope objective. DNO and VNO indicate the dorsal or ventral side of the embryo near the objective, respectively. (C) Cellular GFP intensity heatmap comparing experimental replicates of embryos with the identical or distinct orientation. Cells near and far from the center Z plane are shown separated. (D) Differences in cellular GFP intensity in multiple embryos of the identical integration strains with distinct orientations (red for VNO and green for DNO) were used to model the attenuation factor. The upper panel shows the raw result, and the lower panel shows the result adjusted by using an attenuation factor of 0.054. (E) Representative examples showing the performance of attenuation correction. Scatter plots show the cellular GFP expression in embryos of the same integration strain that are with different orientations during imaging before (top) and after (bottom) the compensation of Z attenuation effect. (F) Distribution of the average Pearson correlation coefficient (left) and consistency of binary expression status (right) between cellular GFP expression in experimental replicates. (G) Tree visualization of the fractions of integration sites at which the GFP is expressed in each cell.

**Figure S2.**
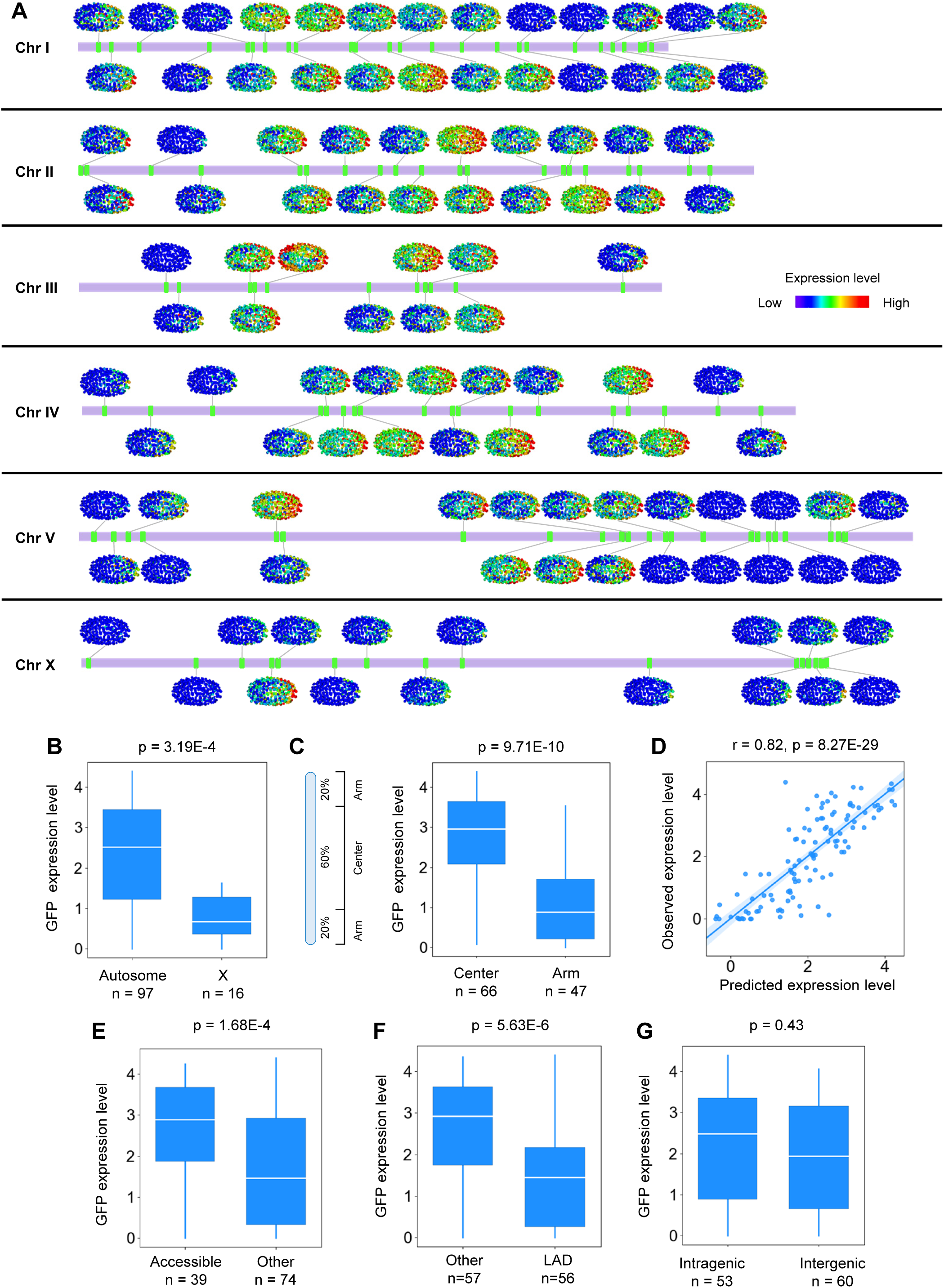
Positional Effects on GFP Expression Reliably Indicate Chromatin Activity Across the Genome, related to Figure 1. (A) Positional effects on GFP expression in 364 identity-resolved cells. Each ellipsoid is a 3D rendering of a standardized embryo, with dots indicating embryonic cells and the color gradient indicating GFP expression level. (B) Comparison of GFP expression levels between autosomes and the X chromosome. (C) Comparison of GFP expression levels between center and arm regions of chromosomes. (D) Correlation of the observed GFP expression level with that predicted by histone modifications. (E) Comparison of GFP expression levels between accessible and non-accessible chromatin. (F) Comparison of GFP expression levels between LAD and non-LAD regions. (G) Comparison of GFP expression between intragenic and intergenic regions. All inter-group statistics were performed by the Mann-Whitney U test; correlation statistics were performed by Pearson correlation.

**Figure S3.**
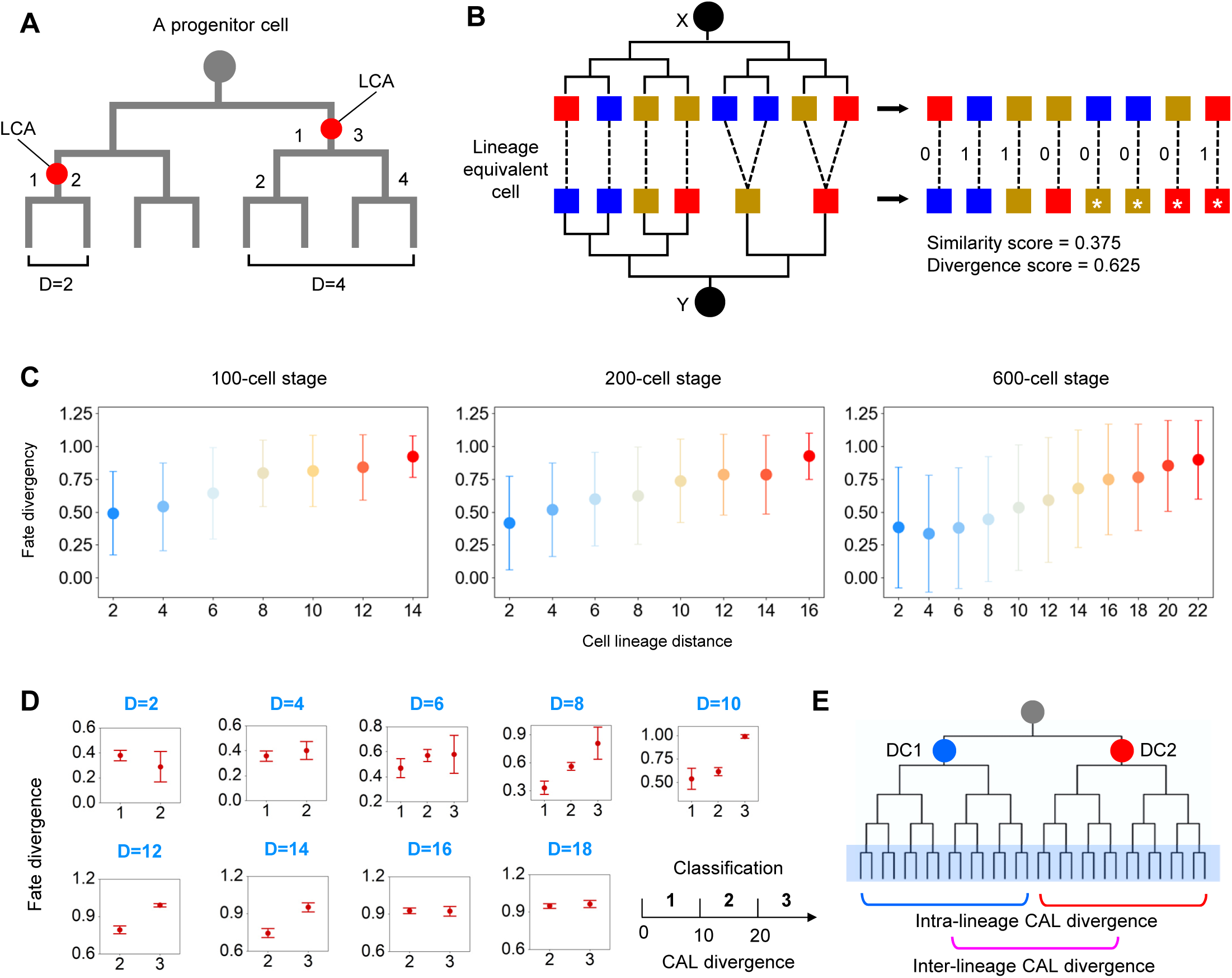
Dynamics of CAL Across Cell Lineages, related to Figure 2. (A) Definition of cell lineage distance. Calculation of cell lineage distance is shown for two examples in which the red circle indicates the lowest common ancestor (LCA) of the two target cells and the numbers list all cell divisions that lead to the two cells from the LCA. (B) Schematic comparing the developmental fate of progenitor cells. The developmental fate of a progenitor cell (circle) is represented as the combinatorial pattern of tissue types (colored squares) it gives rise to following the cell lineage tree. Fate divergence between two progenitor cells (x and y) was quantified by first determining whether the tissue fates are equal (score = 1) or not (score = 0) in each lineal equivalent terminal cell that was then averaged. In case two progenitor cells produced different numbers of terminal cells, the smaller lineage (y) was expanded to match the larger (x) one, cells and tissue fates are expanded (stars) in the corresponding lineage branches. (C) Changes in fate divergence following the increase of cell lineage distance at 100-, 200-, and 600-cell stages. (D) Each panel compares changes in fate divergence between cells with different levels of CAL divergence (divided into three bins). (E) Strategy used to identify CAL transition points. For each pair of daughter cells (DC1 and DC2) produced by a progenitor cell (gray circle), intra- (blue and red lines) and inter-daughter-lineage CAL divergences (magenta line) were compared between terminal cells produced by the two daughter cells to determine whether the mother cell is a CAL transition point.

**Figure S4.**
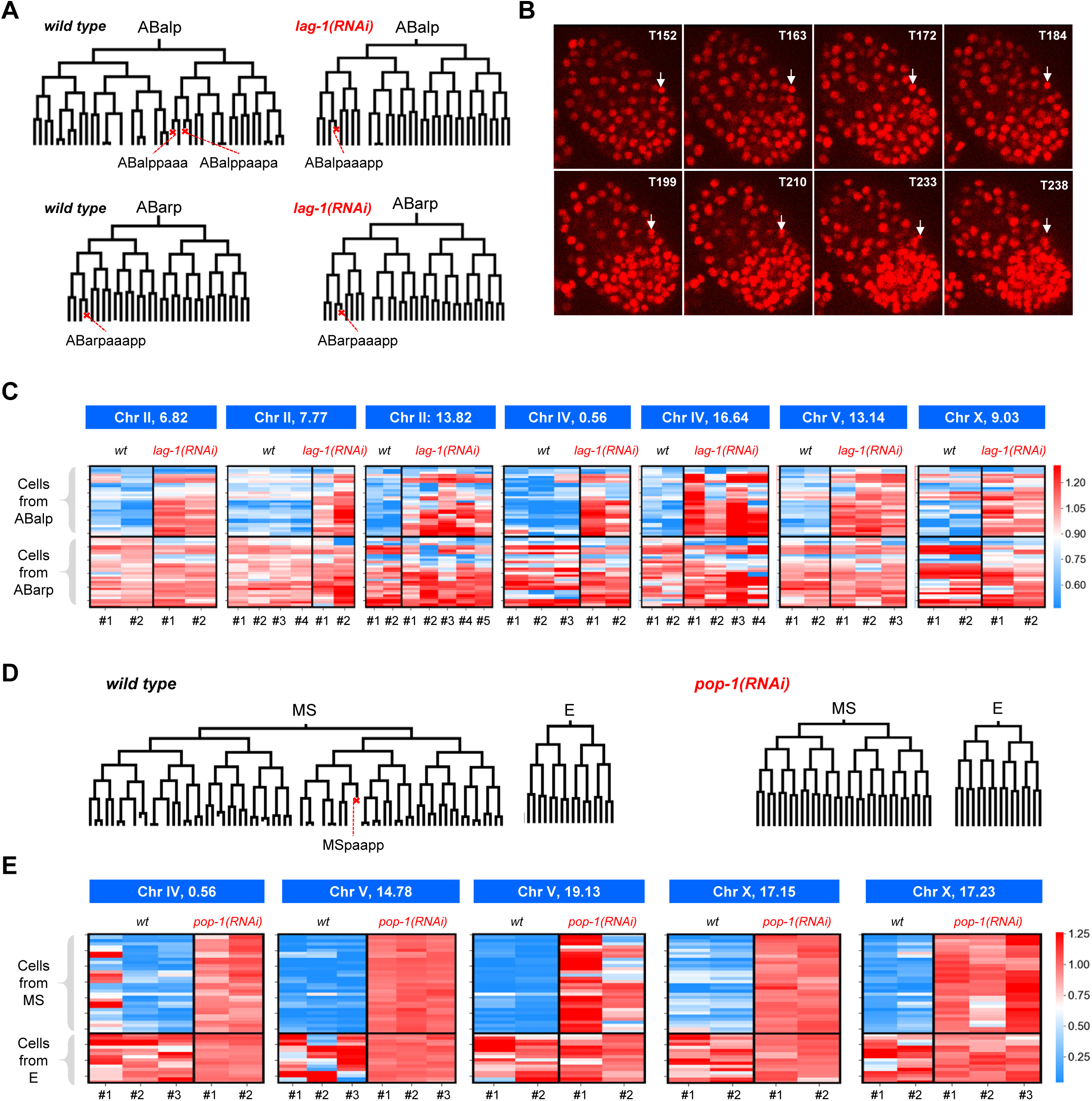
CAL is Coupled to Lineage Fate, related to Figure 3. (A) Cell lineage trees and characteristic programmed cell death (red cross) of the ABalp and ABara lineages in wild-type and *lag-1(RNAi)* embryos. (B) A characteristic sequence of morphological changes in the nucleus was used to determine whether a cell undergoes programmed cell death. (C) Each heatmap shows the relative expression level of GFP integrated into a specific position (indicated above) in cells from the ABalp and ABara lineages in the wild-type and *lag-1(RNAi)* embryos. Cellular GFP expression levels were normalized to the mean value of cells from the ABarp lineage for both genotypes. (D) Cell lineages and characteristic programmed cell death (red cross) in the EMS lineage of wild-type and *pop-1(RNAi)* embryos. (E) Each heatmap shows the relative expression level of GFP integrated into a specific position (indicated above) in cells from the MS and E lineages in the wild-type and *pop-1(RNAi)* embryos. Cellular GFP expression levels were normalized to the mean value of cells from the E lineage in both genotypes.

**Figure S5.**
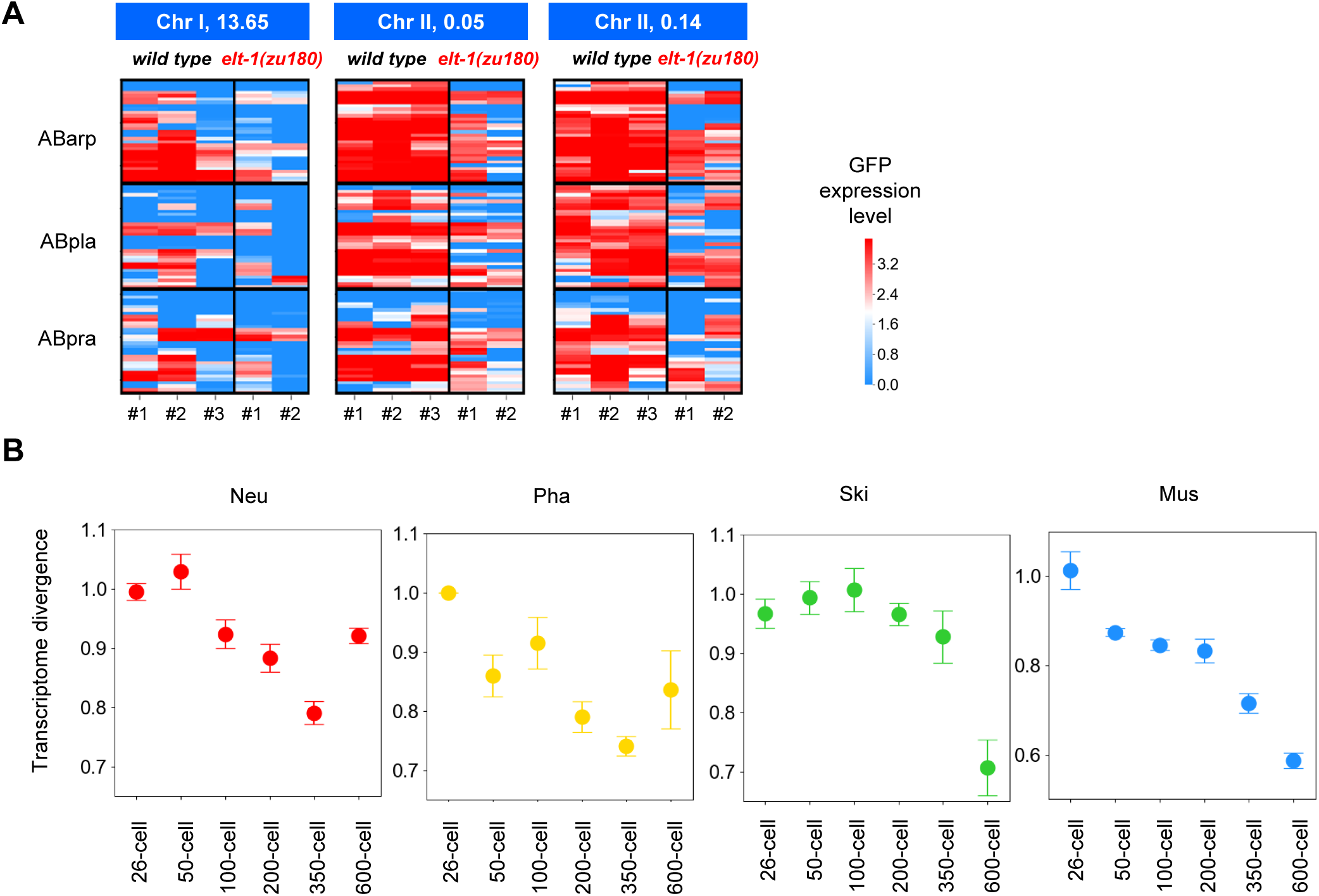
Dynamics and Implications of CAL During Tissue Fate Differentiation, related to Figure 4. (A) Each heatmap shows the expression level of GFP integrated into a specific position (indicated above) in cells from the ABarp, ABpla, and ABpra lineages that normally differentiate into skin in all analyzed wild-type and *elt-1(zu180)* embryos. (B) Transcriptome divergence between cells destined to differentiate into the same tissue type at different embryonic stages. The intestine was not included because the lineage identities of individual cells were highly indistinguishable. Data are presented as mean ± 95% CI.

**Figure S6.**
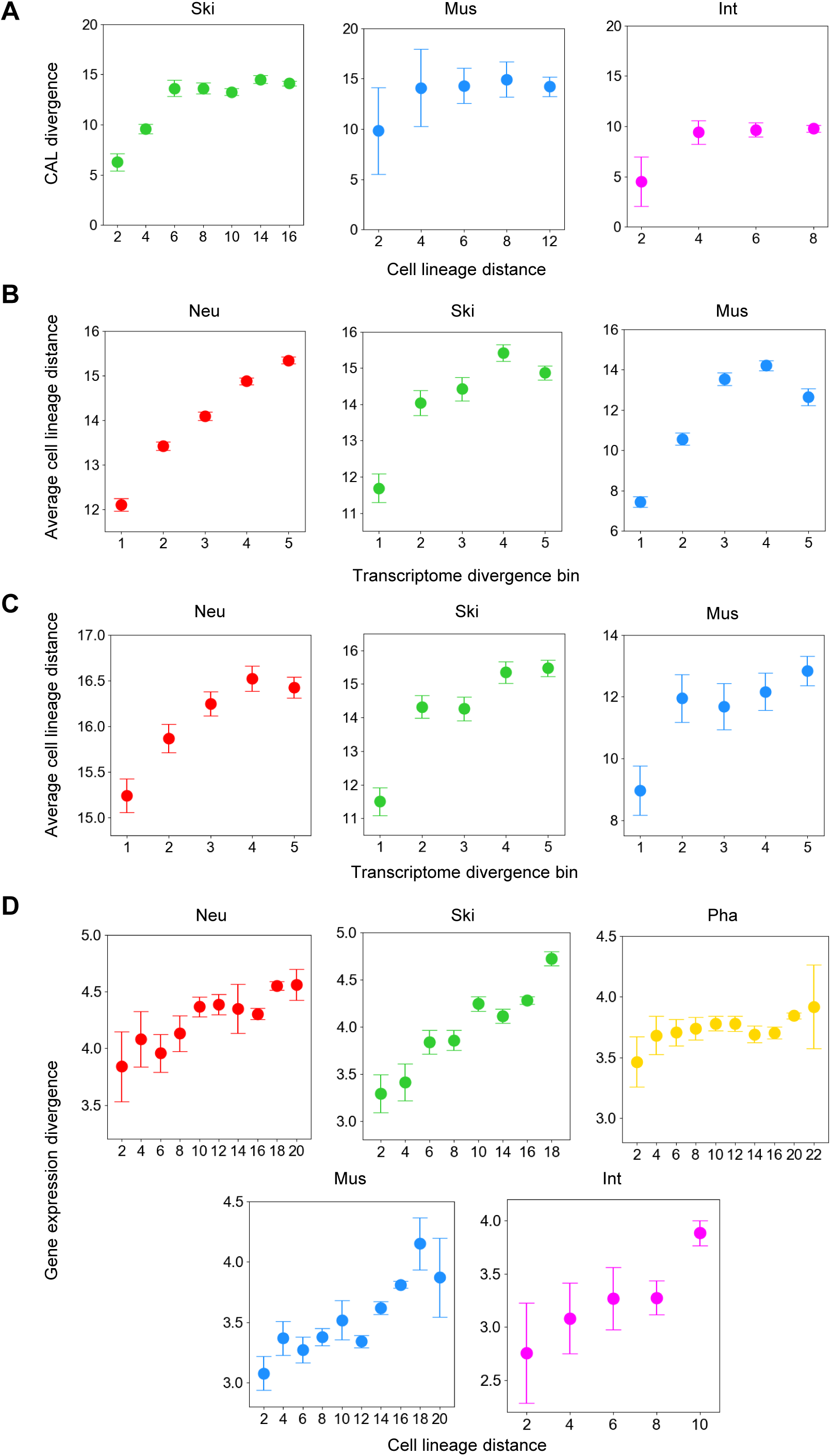
Lineage-Origin-Dependent Intra-Tissue Heterogeneity on CAL and Gene Expression, related to Figure 5. (A) Changes in CAL divergence following an increase in cell lineage distance between cells for all post-mitotic intra-tissue cells. The neuronal system and pharynx were not included due to a small number of post-mitotic cells in the 350-cell stage. Data are presented as mean ± 95% CI. (B-C) Comparison of average cell lineage distance between cells at the 350-cell (B) and 600-cell (C) stages with different transcriptome divergences. In many cases, two possible lineage identities were assigned to a cell; the classification of cells based on lineage distance is inaccurate. We therefore classified transcriptome divergence between cells evenly into five bins and compared the average cell lineage distance between cells in each bin. The pharynx and intestine were not included because many of the cells were assigned more than two possible lineage identities, which would compromise the accuracy of the results. Data are presented as mean ± 95% CI. (D) Changes in gene expression divergences following the increase of cell lineage distance between cells in the L1 larvae stage. Data are presented as mean ± 95% CI.

**Figure S7.**
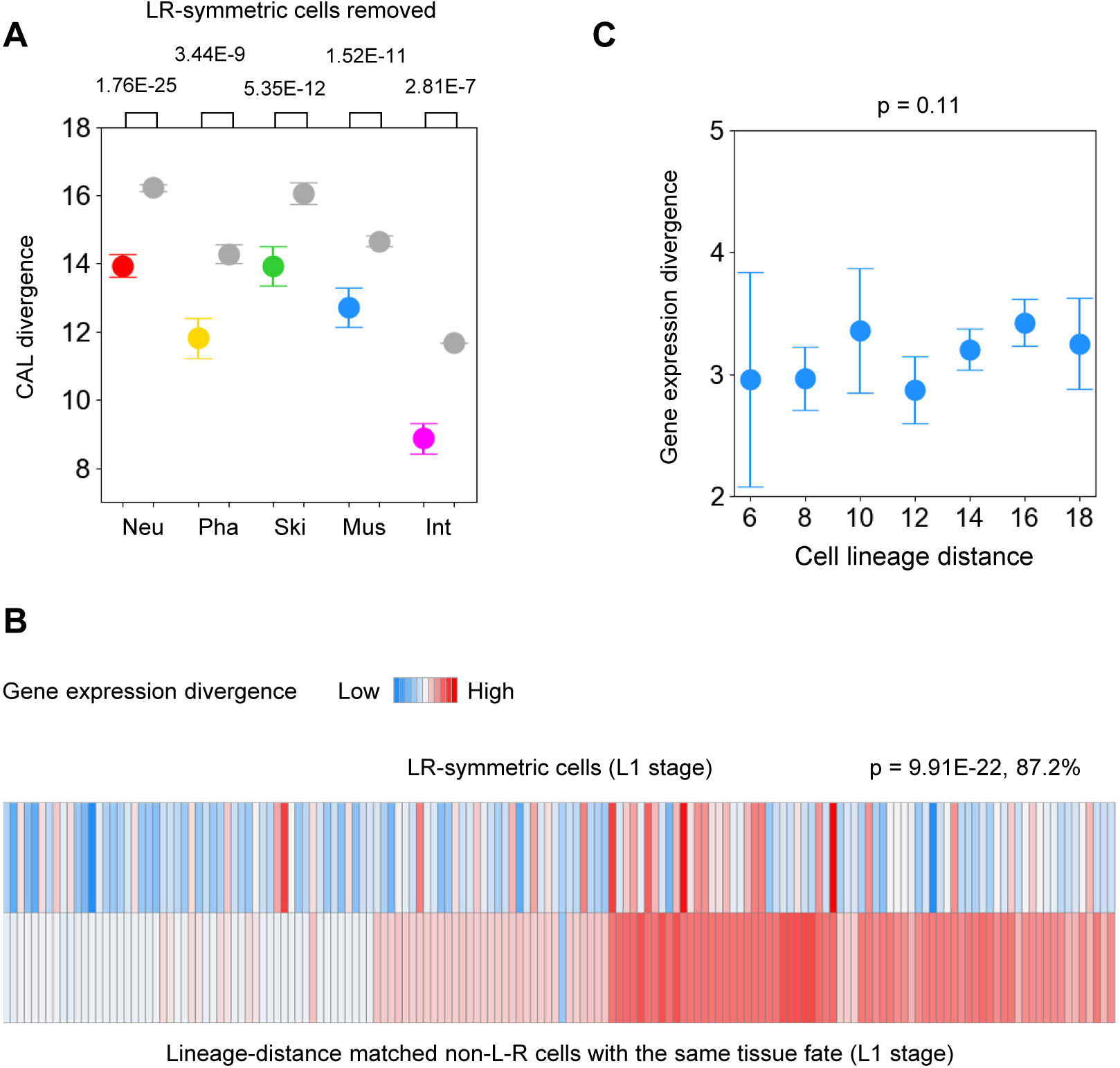
Predetermination of Regulatory States During L-R Symmetry Establishment, Related to Figure 6. (A) Comparison of CAL divergence between intra-tissue cells and lineage-distance-matched inter-tissue cells after removing the L-R symmetric cells from intra-tissue cells. Data are presented as mean ± 95% CI. Statistics: Mann-Whitney U test. (B) Heatmap visualization of gene expression divergence in L1 stage larvae between left and right cells in each pair of L-R symmetric cells/progenitors (top) and between lineage distance matched non-L-R symmetric cells from the same tissue type (bottom). Statistics: Wilcoxon signed-rank test, n = 156. (C) Gene expression divergence between L-R symmetric cells with different cell lineage distances. Data are presented as mean ± 95% CI.

### 7. SUPPLEMENTARY TABLE LEGENDS

Table S1. *C. elegans* Strains, related to STAR Methods

Names and genotypes of the *C. elegans* strains used in this study.

**Table S2. Quantification of Position-Effects on GFP Expression in Single Embryonic Cells to Indicate Chromatin Activity,** related to Figure 1

Sheet 1: Chromosomal locations of 113 integration strains used to assay chromatin activity. Sheet 2: Expression levels of GFP at 113 genomic positions (268 embryos) in 722 traced embryonic cells. Ter indicates the traced terminal cells at the 350-cell stage and Pro indicates earlier intermediate progenitor cells that generate the terminal cells. Sheet 3: CAL of 364 traced terminal somatic cells. Sheet 3: Correlation between CAL and endogenous gene expression in identity matched cells. Sheet 4: Mean expression levels of GFP at each integration site and related genetic and epigenetic features. Sheet 5: Correlation between expression levels of GFP to endogenous genes in a 500-kb interval centered on the integration sites in identity match cells.

**Table S3. Dynamics of CAL Across Cells from Different Lineages,** related to Figure 2

Sheet 1: Mean CAL divergence and fate divergence of each cell compared to all other cells as a function of cell lineage distance. Sheet 2: CAL transition points and associated statistics.

**Table S4. Dynamics of CAL Across Cells from Different Tissues,** related to Figure 4

Sheet 1: Tissue types and differentiation state of 364 traced terminal somatic cells. Sheet 2: CAL divergence between intra- and inter-tissue cells of each tissue type.

**Table S5. Lineage Origins Affects Intra-Tissue CAL Heterogeneity**, related to Figure 5

Sheet 1: CAL divergence between intra- and inter-tissue cells at different cell lineage distances. Sheet 2: Cell clusters.

**Table S6. CAL and Gene Expression Divergence Between L-R Symmetric Cells,** related to Figure 6

Sheet 1: CAL divergence between L-R symmetric cells/progenitor cells. Sheet 2: Gene expression divergence between L-R symmetric cells in the L1 stage. Sheet 3: Comparison of CAL divergence between differentiated L-R symmetric cells and between their mother cells.

**Table S7. CDCA regions Predict Structural and Functional Relevance of the Genome,** related to Figure 7

Sheet 1: Pair-wise divergence (Euclidean distance) of inter-cell dynamics of chromatin activity between 113 genomic positions. Sheet 2: Clusters of CDCA regions. Sheet 3: relative frequency of CDCA regions in TAD and non-TAD regions. Sheet 4-7: enrichment of functionally related genes near CDCA regions.

## Notes

### Competing Interest Statement

The authors have declared no competing interest.

